# Curing autoimmune diabetes with islet and hematopoietic cell transplantation after CD117 antibody-based conditioning

**DOI:** 10.1101/2025.09.05.673576

**Authors:** Preksha Bhagchandani, Stephan A. Ramos, Bianca Rodriguez, Xueying Gu, Shiva Pathak, Yuqi Zhou, Yujin Moon, Nadia Nourin, Charles A. Chang, Jessica Poyser, Brenda J. Velasco, Weichen Zhao, Hye-Sook Kwon, Richard Rodriguez, Diego Burgos, Mario A. Miranda, Everett Meyer, Judith A. Shizuru, Seung K. Kim

## Abstract

Mixed hematopoietic chimerism after allogeneic hematopoietic cell transplantation (HCT) promotes tolerance of transplanted donor-matched solid organs, corrects autoimmunity, and could transform therapeutic strategies for autoimmune type 1 diabetes (T1D). However, development of non-toxic bone marrow conditioning protocols is needed to expand clinical use. We developed a chemotherapy-free, non-myeloablative (NMA) conditioning regimen that achieves mixed chimerism and allograft tolerance across MHC barriers in NOD mice. We obtained durable mixed hematopoietic chimerism in prediabetic NOD mice using anti-c-Kit monoclonal antibody, T-cell depleting antibodies, JAK1/2 inhibition, and low-dose total body irradiation prior to transplantation of MHC-mismatched B6 hematopoietic cells, preventing diabetes in 100% of chimeric NOD:B6 mice. In overtly diabetic NOD mice, NMA conditioning followed by combined B6 HCT and islet transplantation durably corrected diabetes in 100% of chimeric mice without chronic immunosuppression or graft-versus-host disease (GVHD). Chimeric mice remained immunocompetent, as assessed by blood count recovery and rejection of 3^rd^ party allogeneic islets. Adoptive transfer studies and analysis of autoreactive T cells confirmed correction of autoimmunity. Analysis of chimeric NOD mice revealed central thymic deletion and peripheral tolerance mechanisms. Thus, with NMA conditioning and cell transplantation, we achieved durable hematopoietic chimerism without GVHD, promoted islet allograft tolerance, and reversed established T1D.

**Graphical Abstract:** 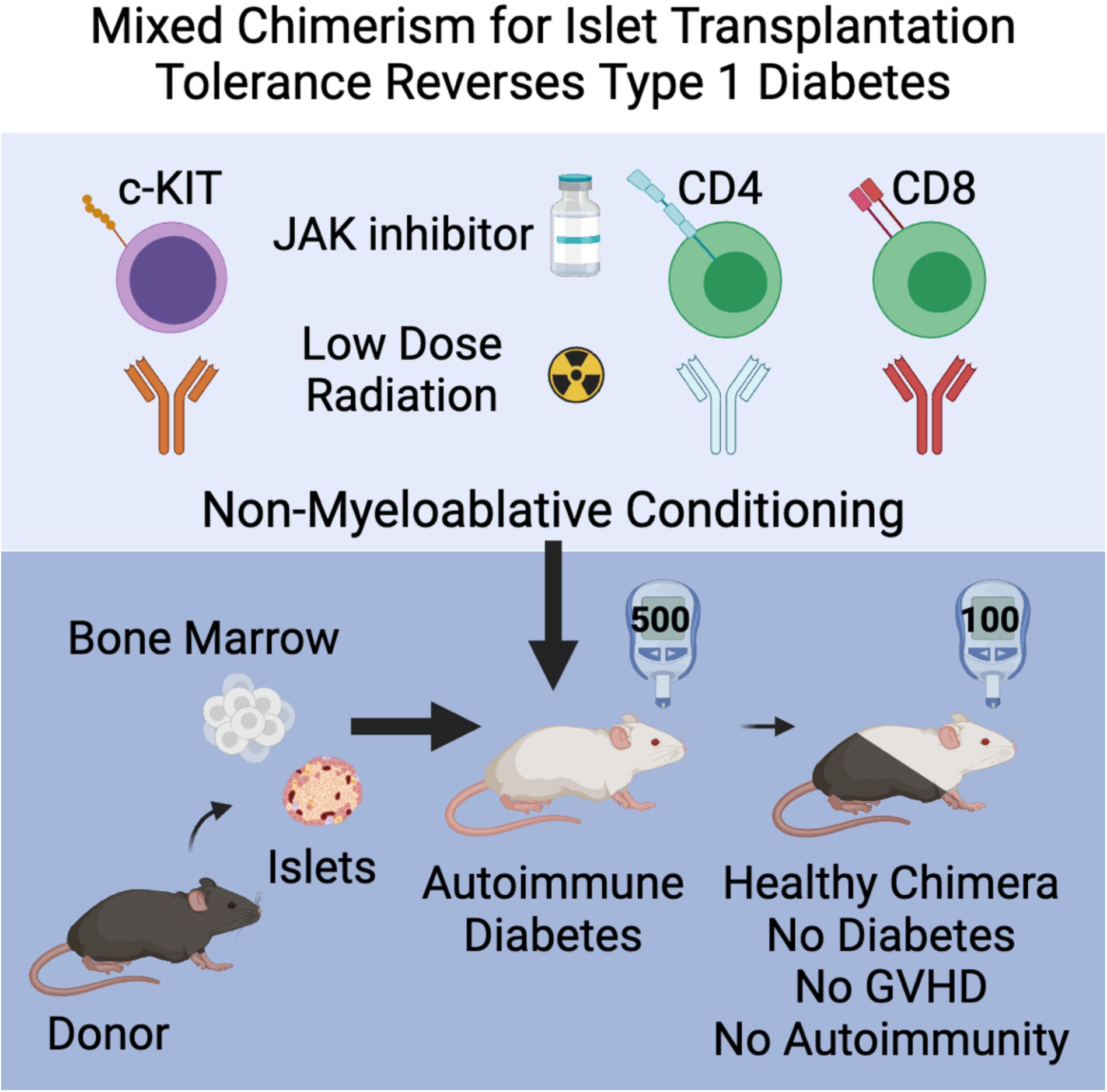

## INTRODUCTION

Type 1 diabetes (T1D) results from autoimmune destruction of insulin-producing pancreatic islet β cells, resulting in life-long dependence on exogenous insulin (1). Islet transplantation from cadaveric human donor pancreas is a promising, FDA-approved treatment for T1D (2, 3). However, clinical islet transplantation is complicated by two immunological obstacles: (**1**) allogeneic rejection of MHC-mismatched tissues and (**2**) recurrent islet-specific autoimmune rejection (4). The current clinical standard in islet transplantation includes use of life-long systemic immunosuppressive drugs, often associated with adverse effects such as β cell toxicity, malignancy, risk of infection, and chronic rejection. These risks preclude adoption of islet transplantation for most T1D patients, challenging physicians and scientists to improve strategies for promoting immune tolerance instead of systemic immunosuppression.

Mixed hematopoietic chimerism after donor hematopoietic cell transplantation (HCT), allows for donor-matched solid organ tolerance (5–8). Furthermore, allogeneic HCT has been shown to correct autoimmunity in mice and human (9), supporting the view that autoimmune diseases like T1D reflect defects in hematopoietic stem cells (HSCs). Mixed hematopoietic chimerism, where donor and host HSCs co-exist, is achieved by ‘conditioning’ the recipient with a regimen that creates bone marrow niche space and transiently lowers immunological barriers to donor HSC transplantation and engraftment (10). Unlike in cancer therapies, where HCT follows high-dose radiation therapy (XRT) and/or chemotherapy-based conditioning, immunological tolerance promoted by mixed chimerism for non-malignant conditions, like T1D, mandates safer non-myeloablative (NMA) conditioning protocols. Thus, the development and improvement of NMA conditioning protocols to promote durable mixed chimerism could advance strategies for solid organ transplant tolerance (11). For T1D, it will also be essential to evaluate the efficacy of NMA conditioning protocols, mixed chimerism and islet transplantation in animals with established disease. Findings from pre-clinical studies of mixed chimerism and islet transplantation tolerance in autoimmune diabetes have precluded clinical adoption due to risks of graft versus host disease (GVHD), and use of conditioning regimes and reagents not clinically portable, including conditioning chemotherapy or intensive XRT (12–17).

We have built on our previous work with anti-CD117-based conditioning followed by HCT and islet transplantation to reverse diabetes in a non-autoimmune mouse model (18–20), to generate a novel, non-toxic conditioning regimen that promotes mixed chimerism across MHC barriers in NOD mice with autoimmune diabetes. This NMA regimen includes (**1**) antibody targeting of CD117, a receptor tyrosine kinase essential for HSC proliferation and survival through the binding of stem cell factor (21), and (**2**) inhibition of JAK-STAT signaling to overcome substantial NK cell and T cell barriers to allogeneic transplant in the setting of autoimmunity (22–24). With this conditioning strategy, we achieved durable allogeneic islet transplantation tolerance and autoimmune diabetes reversal in NOD mice. These findings in T1D using clinically portable reagents provide proof of concept for safer bone marrow and curative islet transplantation, without GVHD.

## RESULTS

### Non-myeloablative conditioning promotes robust and durable allogeneic donor chimerism

NOD mice are resistant to the immunosuppressive effects of radiation, making them challenging to engraft with HSCs (25). Previously, we achieved durable hematopoietic chimerism after allogeneic HCT in B6 mice using non-myeloablative (NMA) conditioning with 300 cGy total body irradiation (TBI), αCD117 antibody, and transient T-cell depletion (TCD) with αCD4 and αCD8 antibodies (18). However, after conditioning 8-week-old prediabetic NOD mice with this regimen, followed by transplantation with allogeneic B6 donor bone marrow (Figure 1A and Supplemental Figure 1A), we did not achieve long-term multilineage engraftment (Supplemental Figure 1B). Analysis of NOD peripheral blood and lymphoid organs suggested that host NK and T cells might be insufficiently suppressed by that conditioning regimen (Supplemental Figure 1C). To test this hypothesis, we added a daily dose of baricitinib, a JAK1/2 inhibitor that targets NK cells and T cells found to be well-tolerated in pre-clinical and clinical studies of allogeneic HSC engraftment in non-diabetic hosts (12 days total; Figure 1B and Supplemental Figure 1C) (22, 26, 27). Prediabetic NOD mice were conditioned from day −4 to day +8 with 225 or 250 cGy TBI on day −2, transplanted with whole bone marrow (WBM) on day 0, and followed for up to 20 weeks (Figure 1B). Analysis at 4 weeks after HCT revealed robust donor chimerism in all peripheral blood lineages regardless of radiation dose (Overall: 83.8 ± 5.9%, CD3^+^: 76.6 ± 3.8%, CD19^+^: 91.3 ± 2.2%, CD11b^+^: 81.7 ± 6.6% CD49b^+^: 79.6 ± 6.4%; *n* = 15; Figure 1C). Stable mixed chimerism was maintained in peripheral blood throughout the 20-week experiment period in 14 of 15 recipients, hereafter referred to as “prediabetic NOD:B6” (Figure 1D). Similarly, endpoint analysis of the spleen and bone marrow showed high chimerism levels (Figure 1, E and F). We also confirmed engraftment of donor Lin^−^Sca1^+^cKit^+^ (LSK) HSCs and persistence of host LSK HSCs (Donor LSK chimerism: 61.5 ± 6.5%; Figure 1F). Using lineage-depleted hematopoietic stem and progenitor cells (HSPCs) instead of WBM, we achieved similar outcomes (Supplemental Figure 1D). Furthermore, we established a ‘subminimal’ radiation dose at which we no longer achieve engraftment (200 cGy; Supplemental Figure 1E). Thus, we generated durable mixed hematopoietic chimerism in prediabetic NOD mice with a NMA regimen combining low-dose TBI with αCD117 and baricitinib.

**Figure 1.**
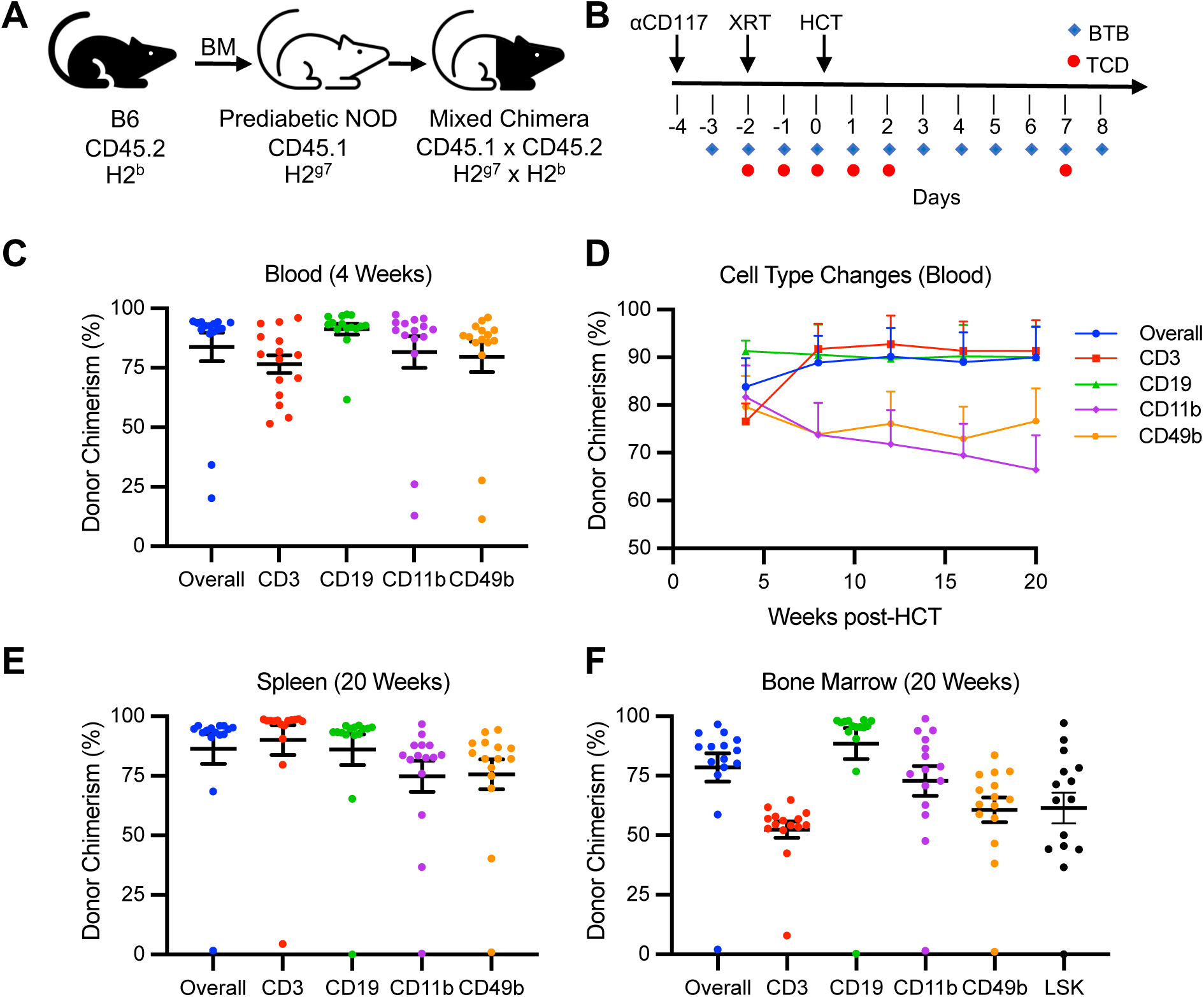
Non-myeloablative conditioning promotes robust and durable allogeneic donor chimerism in a radioresistant NOD model. **(A)** Transplantation schematic and strains used. **(B)** Reduced intensity conditioning regimen. **(C)** Chimerism analysis of peripheral blood 4 weeks after conditioning and HCT depicting overall, CD3^+^ T cell, CD19^+^ B cell, CD11b^+^ myeloid cell, and CD49b^+^ NK cell donor (CD45.2^+^) chimerism (*n* = 15, sum of 3 independent experiments). **(D)** Longitudinal chimerism analysis of peripheral blood over a 20-week period combined from 3 separate experimental cohorts (*n* = 5 per cohort). **(E)** Chimerism analysis of host spleen at 20 weeks post-HCT (*n* = 15, sum of 3 independent experiments). **(F)** Chimerism analysis of host bone marrow at 20 weeks post-HCT including Lin^−^Sca1^+^cKit^+^ (LSK) HSCs (*n* = 15, sum of 3 independent experiments). C-F. Data are represented with mean ± SEM. XRT = radiation therapy; HCT = hematopoietic cell transplant; BTB = baricitinib; TCD = T cell depletion

After WBM transplantation, we did not observe clinical signs of GVHD (e.g., mucosal changes, skin rash, or weight loss), or detect evidence of intestinal infiltration (Supplemental Figure 2, A and B). Longitudinal body weight measures showed excellent weight gain from HCT to experimental endpoint (39.0 ± 2.1%; Supplemental Figure 2A). Furthermore, complete and differential blood cell counts (CBCs) showed good recovery at 4 weeks after HCT, including white blood cell counts (WBC) and immune cell subsets, without the need for additional supportive care, such as blood transfusion, indicating healthy reconstitution of bone marrow and HSC function (Supplemental Figure 2, C and D). Thus, after NMA conditioning, multiple indices showed that chimeric NOD:B6 mice had good functional status and health, with hematopoietic recovery and no GVHD.

### Mixed chimerism prevents autoimmune diabetes and establishes donor-specific islet tolerance

Hematopoietic chimerism can prevent and correct autoimmunity (9, 28). Thus, we asked if mixed chimerism after αCD117-based NMA conditioning could prevent autoimmune diabetes in NOD mice. Although naïve prediabetic NOD mice at 8 weeks of age are normoglycemic, they exhibit insulitis by pancreatic islet histology with predominant B and T cell infiltrate (Supplemental Figure 3, A and B). Between 14-24 weeks of age, 60% of NOD mice in our colony developed diabetes, (*n* = 9/15; Figure 2, A and B). In conditioned NOD mice *without* HCT (‘conditioned controls’), we observed delayed diabetes onset, beginning 16 weeks after conditioning, reduced diabetes incidence to 30%, and insulitis on pancreatic histology (*n* = 3/10; Figure 2, A-C). These features, including timing of diabetes onset, likely reflect the delayed recovery of bone marrow and immune function after conditioning NOD mice without subsequent HCT. Insulitis scoring of pancreatic histology revealed a similar proportion of islets with insulitis (70-75%) in both naïve NOD and conditioned controls, despite delayed progression and reduced incidence (Figure 2C), and no difference in infiltrate composition by cell type (Supplemental Figure 3B). By contrast, NMA conditioning *and* HCT prevented diabetes onset in 100% of NOD:B6 (*n* = 19/19; Figure 2, A and B). Pancreatic histology in chimeric NOD:B6 mice at 20 weeks post-HCT showed a majority of islets without CD45^+^ immune cells in the peri-islet region and few islets with minimal peri-insulitis (Figure 2C), similar to B6 mice (Supplemental Figure 3A). Insulitis scoring of pancreatic histology revealed less than 15% of islets with peri-insulitis (Figure 2C). Analysis of those peri-islet immune cells revealed most to be of donor (CD45.2^+^) origin (Figure 2D), possibly indicating cells that suppressed autoimmunity (29). The majority of these peri-islet cells were B and T cells along with few myeloid cells (Figure 2D). Thus, mixed hematopoietic chimerism after αCD117-based NMA conditioning corrected autoimmunity and prevented diabetes onset and insulitis in prediabetic NOD mice.

**Figure 2.**
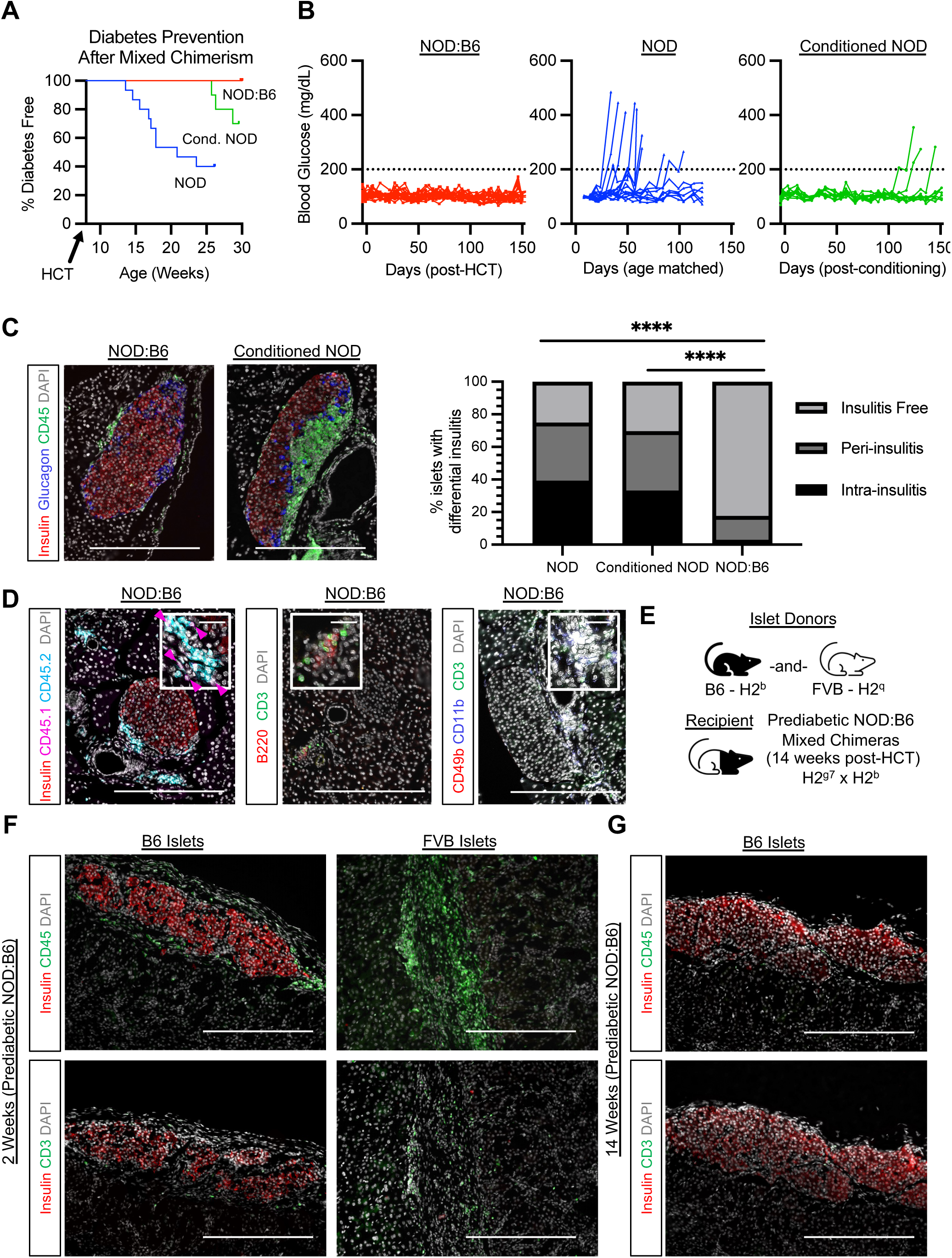
Mixed chimerism prevents autoimmune diabetes and establishes donor-specific islet tolerance. **(A)** Autoimmune diabetes development curves of prediabetic NOD:B6 mixed chimeric mice (*n* = 19 from 4 independent experiments), conditioned controls (*n* = 10), and naïve NOD (*n* = 15). *P* < 0.0001 for chimeras vs. naïve NOD and *P* < 0.01 for chimeras vs. conditioned controls using log-rank (Mantel-Cox) test. *P* < 0.0001 for chimeras vs. naïve NOD and *P* < 0.05 for chimeras vs. conditioned controls using Fisher’s exact test. *P* < 0.05 (Mantel-Cox) and no significance (ns, Fisher’s exact) for conditioned controls vs. naïve NOD. **(B)** Non-fasting blood glucose of NOD:B6 chimeras (red) compared to naïve NOD (blue) and conditioned controls (green). Dotted line indicates normoglycemic threshold (200 mg/dL). **(C)** Percentage of different stages of insulitis in individual islets and representative histology of pancreas from NOD:B6 20 weeks post-HCT (*n* = 14) compared to naïve NOD (*n* = 5) and conditioned controls (*n* = 10). Statistical analyses done using ξ^­^test. **(D)** Representative pancreatic histology of NOD:B6 chimeras at 20 weeks post-HCT stained for insulin, CD45.1 (host), and CD45.2 (donor) or B220 and CD3 or CD49b, CD11b and CD3 (*n* = 3-6). Pink arrows indicate CD45.1^+^ immune cells. Inset scale bars 20µm. **(E)** Experimental transplantation schematic. Prediabetic NOD:B6 mixed chimeras received B6 islets in the left kidney and FVB islets in the opposite kidney at 14 weeks post-HCT. **(F)** FVB and B6 islet grafts 2 weeks after islet transplantation in mixed chimeras stained for insulin and CD3 or CD45 (*n* = 3). **(G)** B6 islet grafts 14 weeks after islet transplantation in mixed chimeras stained for insulin and CD3 or CD45 (*n* = 4). D-G. Scale bars = 200μm unless otherwise specified. ****P < 0.0001 HCT = hematopoietic cell transplantation.

To test if mixed hematopoietic chimerism in prediabetic NOD mice promoted donor-matched islet allograft and autoantigen tolerance, we transplanted islets from B6 or third-party FVB (H2^q^) sex-matched control mice into prediabetic NOD:B6 chimeras 14 weeks after HCT (Figure 2E). Two weeks after transplantation, recovered B6 islet grafts remained intact in prediabetic NOD:B6 chimeras, with little to no immune infiltration (Figure 2F). By contrast, third-party FVB islet grafts were readily rejected, showing heavy infiltration by immune cells, and contained no detectable INS^+^ cells (Figure 2F). Donor-matched B6 islet grafts assessed up to 14 weeks after islet transplantation also appeared intact, with little or no immune cell infiltrate (Figure 2G). Thus, mixed hematopoietic chimerism in prediabetic NOD prevented autoimmune diabetes, maintained self-tolerance, promoted long-term donor-matched islet allotolerance, and preserved immunocompetence against foreign antigens.

### Curing autoimmune diabetes with allogeneic hematopoietic cell and islet transplantation

To investigate if combined HCT and islet transplantation could reverse overt diabetes in NOD mice (hyperglycemia > 2 weeks), we transplanted allogeneic B6 WBM and islets from either B6 (donor-matched) or FVB (third-party) donors in the subcapsular renal space (Figure 3A). Diabetic mice were maintained on exogenous insulin (Methods) prior to NMA conditioning, which was discontinued on day 0 after simultaneous HCT and islet transplantations (∼400 IEQ). After 4 weeks, multi-lineage mixed chimerism was observed in the peripheral blood of diabetic NOD recipients (Overall: 74.9 ± 6.4%, CD3^+^: 67.5 ± 4.8%, CD19^+^: 80.0 ± 6.8%, CD11b^+^: 59.9 ± 7.6% CD49b^+^: 66.1 ± 7.3%; *n* = 19; Figure 3B). Of nine mice transplanted with 400 IEQ and followed long term, all exhibited durable mixed chimerism through 20 weeks post-transplantation (Figure 3C). This was further confirmed by endpoint analysis of the bone marrow at 20 weeks (donor LSK chimerism: 45.0 ± 8.1%; Figure 3D). In two control NOD:B6 mice, we transplanted ∼200 B6 IEQ, resulting in uncorrected hyperglycemia peri-transplantation. Neither of these mice achieved long-term mixed hematopoietic chimerism; given prior studies indicating that hyperglycemia can alter the bone marrow microenvironment and possibly delay bone marrow engraftment (30–33), this may be an important variable to consider for clinical translation.

**Figure 3.**
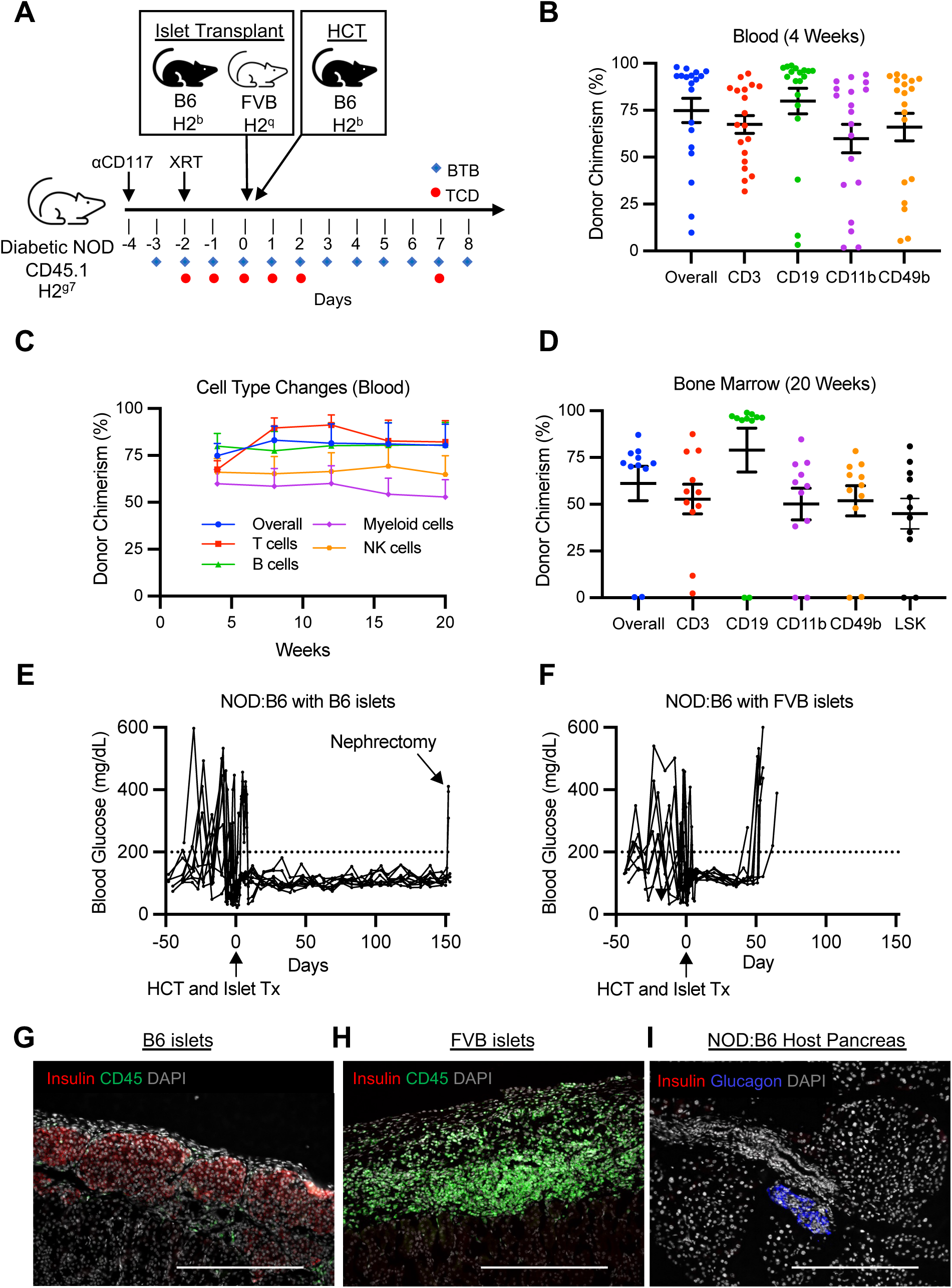
Curing autoimmune diabetes with allogeneic hematopoietic cell and islet transplantation. **(A)** Experimental conditioning and transplantation timeline. **(B)** Multilineage chimerism analysis 4 weeks after HCT (*n* = 19, 6 independent experiments). **(C)** Longitudinal chimerism analysis of peripheral blood over a 20-week period post-HCT (*n* = 11). **(D)** Chimerism levels of immune cell subtypes in the bone marrow of mixed chimeras at 20-weeks after HCT (*n* = 11). **(E)** Non-fasting blood glucose of mice that received B6 islets and developed mixed chimerism (*n* = 9). Arrow indicates nephrectomy (*n* = 3). **(F)** Non-fasting blood glucose of mixed chimeric mice that received FVB islets (*n* = 8). (E, F) Dotted line (200 mg/dL) indicates normoglycemia threshold. **(G)** B6 islet graft 20 weeks after transplantation in NOD:B6 mixed chimera, stained for insulin and CD45 (*n* = 9). **(H)** FVB islet graft 8 weeks after transplantation in NOD:B6 mixed chimera, stained for insulin and CD45 (*n* = 8). **(I)** NOD:B6 host pancreas 20 weeks after transplant stained for insulin and glucagon (*n* = 17). B-E. Data are represented as means ± SEM. Scale bar = 200μm. XRT = radiation therapy; HCT = hematopoietic cell transplant; BTB = baricitinib; TCD = T cell depletion

Diabetic NOD mice with durable mixed hematopoietic chimerism (hereafter, “diabetic NOD:B6”) all exhibited normal blood glucose levels throughout the 20-week follow-up period, with no additional immunosuppression or supplemental insulin required (*n* = 9/9; Figure 3E). To ensure that the donor-matched islet graft was solely responsible for diabetes control, the islet-bearing kidney was removed (“Nephrectomy”), resulting in reversion to diabetes (*n* = 3/3; Figure 3E). Additionally, diabetic conditioned controls without HCT swiftly rejected B6 islets grafts within an average of 39 days after transplant, demonstrating that conditioning alone does not promote tolerance (*n* = 3/3; Supplemental Figure 4, A and B). In contrast to chimeras that received B6 islets, chimeric mice that received third-party FVB islets initially stabilized blood glucose levels but reverted to hyperglycemia in an average of 50 days, indicating rejection of FVB islets, and suggesting that NOD:B6 mice maintained immunocompetence (*n* = 8/8; Figure 3F and Supplemental Figure 4C). Histology of the recovered islet graft at the experimental endpoint revealed intact B6 islets with few CD45^+^ immune cells in NOD:B6 hosts, contrasted by heavy immune cell infiltration and no detectable INS^+^ cells in FVB islet grafts (Figure 3, G and H). Likewise, endogenous pancreatic histology in NOD:B6 mice showed no detectable INS^+^ cells (Figure 3I).

As in prediabetic NOD:B6, we did not observe clinical indices of GVHD. Additionally, longitudinal body weight measures showed good average weight gain from HCT to the experimental endpoint (15.4 ± 2.1%), and complete blood counts indicated healthy hematopoiesis and reconstitution (Supplemental Figure 4, D and E). Thus, multiple indices (glycemia, weight gain, CBCs, immunocompetence) demonstrated good functional status and diabetes reversal in diabetic NOD:B6.

### T cell autoimmunity is corrected in prediabetic and diabetic NOD:B6 chimeras

To assess the mechanisms of autoimmunity correction in NOD:B6 mice, we evaluated the deletion of autoreactive cytotoxic T cells in both prediabetic and diabetic NOD:B6 chimeras. Antigen-specific CD8^+^ T cells reactive to the islet-associated autoantigen glucose-6-phosphatase catalytic subunit-related protein (IGRP) found in both NOD mice and humans with T1D, were measured using MHC Class I (H-2K^d^) IGRP tetramer staining (Methods; Supplemental Figure 5A). Naïve prediabetic NOD and conditioned control NOD mice (20 weeks post-conditioning) show similar levels of IGRP-double-tetramer^+^ CD8^+^ T cells in pancreatic lymph nodes (pancLN) and spleen (Figure 4A). However, these cells were almost completely depleted in prediabetic NOD:B6 mice at 20 weeks post-HCT (Figure 4A). We observed a similar trend in prediabetic NOD:B6 peripheral blood (Supplemental Figure 5B). This depletion was also pronounced in diabetic NOD:B6 mice; we observed nearly complete elimination of IGRP-double-tetramer^+^ CD8^+^ T cells in the pancLN, spleen, and peripheral blood. In contrast, diabetic NOD controls exhibited a marked increase in autoreactive T cells (Figure 4B).

**Figure 4.**
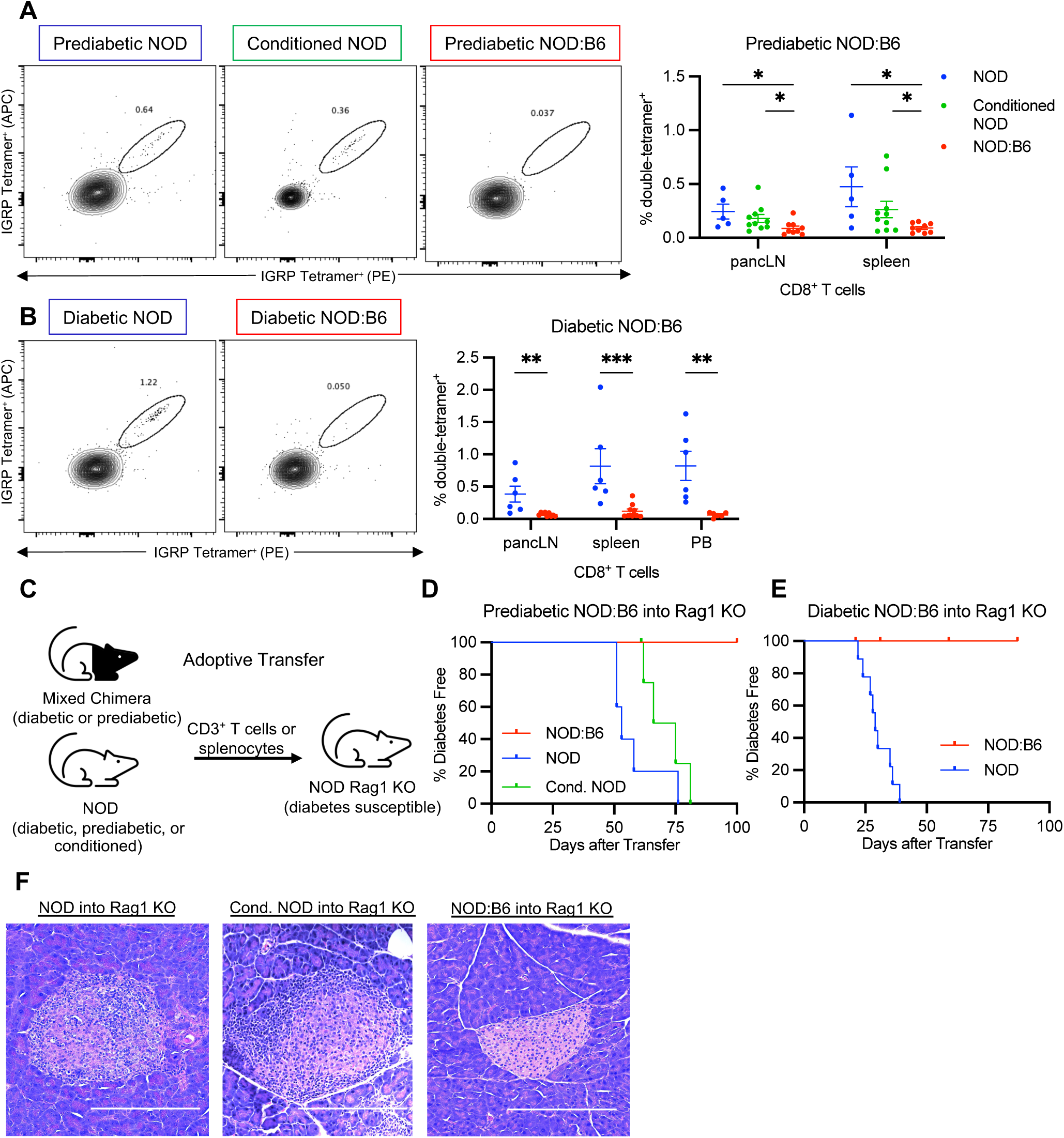
T cell autoimmunity is corrected in prediabetic and diabetic chimeras. **(A)** Representative flow plots and IGRP-double-tetramer^+^ autoreactive cell frequency among CD8^+^ T cells in pancreatic lymph nodes (pancLN) and spleens of prediabetic NOD, conditioned NOD controls, and prediabetic NOD:B6 at 20 weeks post-HCT (*n* = 5-10). **(B)** Representative flow plots and mean ± SEM IGRP-double-tetramer^+^ autoreactive cell frequency in CD8^+^ T cells in the pancLNs, spleens, and peripheral blood (PB) of diabetic NOD and diabetic NOD:B6 20 weeks post-HCT and islet transplantation (*n* = 6-9). **(C)** Adoptive transfer schematic. Diabetes incidence in NOD Rag1 KO mice injected with **(D)** T cells from prediabetic NOD (naïve or conditioned) vs. prediabetic NOD:B6 mice (*P* = 0.0007 and *P* = .0011, respectively, *n* = 5-6), or **(E)** Diabetic NOD vs. diabetic NOD:B6 splenocytes (*P* < 0.0001, *n* = 9). Censored events are unrelated to diabetes (*n* = 1; *n* = 3). Statistical analyses done using Mantel-Cox test. **(F)** Representative H&E staining of NOD Rag1 KO pancreatic islets after adoptive transfer of NOD (*n* = 15), conditioned NOD (*n* = 5), or NOD:B6 (*n* = 12) cells. Scale bar = 200μm. (A-C) Data presented as mean ± SEM. \**P* < 0.05, \*\**P* < 0.01.

Adoptive immune cell transfer into an immunodeficient NOD Rag1 KO mice can assess the autoreactive and diabetogenic potential of T cells from prediabetic and diabetic NOD mice; the recipient NOD Rag1 KO mice are deficient in endogenous T and B cells and susceptible to autoimmune diabetes (34). We used adoptive transfer to assess correction of autoimmunity in NOD:B6 mice (Figure 4C). 2×10^6^ MACS-sorted CD3^+^ T cells from spleens of naïve 12-week-old prediabetic NOD mice, conditioned prediabetic NOD (12 weeks post-conditioning), or prediabetic NOD:B6 chimeras (45% donor T cell chimerism; 12 weeks post-HCT) were transferred into normoglycemic naïve 7-8-week-old NOD Rag1 KO female mice and monitored for diabetes incidence. Prediabetic and diabetic NOD mice transferred autoimmunity to 100% of NOD Rag1 KO recipients (*n* = 6/6), with diabetes developing between 50 to 75 or 25 to 40 days after transfer, respectively (Figure 4, D and E). Conditioned prediabetic NOD controls also transferred autoimmunity to 100% of NOD Rag1 KO recipients followed long-term (*n* = 4/4), with diabetes developing 60 to 80 days after transfer. By contrast, cells from prediabetic NOD:B6 or diabetic NOD:B6 mice did not transfer diabetes, with 100% of the recipient mice remaining diabetes free for up to 100 days (*n* = 6/6 for each, Figure 4, D and E). Pancreatic histology showed marked insulitis, after transfer of T cells from naïve NOD or conditioned NOD donors into Rag1 KO recipients (Figure 4F). In comparison, we observed healthy islet morphology in the pancreas, free of insulitis, after adoptive T cell transfer from NOD:B6 chimeric mice (Figure 4F). Thus, we conclude that islet autoimmunity and diabetogenic potential were eliminated after establishing mixed hematopoietic chimerism in prediabetic and diabetic NOD mice.

### Donor thymic cells are associated with central deletion and thymic Treg development

Tolerance to auto- and allo-antigens in mixed chimerism is primarily established through the education of developing thymocytes to donor and self-antigen via donor antigen presenting cells (APCs) (11). We confirmed the presence of donor B6 APC subtypes known to participate in negative selection, including CD11c^+^ DC subsets, thymus-resident dendritic cells (tDCs; CD8^+^ Sirpα^−^), migratory DCs (mDCs; CD8^−^ Sirpα^+^), and plasmacytoid DCs (pDCs; PDCA-1^+^ B220^+^), as well as B cells (Figure 5A) (35–38). mDCs and pDCs participate in clonal deletion of reactive T cells by transporting peripheral (i.e. donor) antigens to the thymus (39) and both subsets were enriched in prediabetic mixed chimeras compared to conditioned NOD and naïve donor controls (Figure 5B; Supplemental Figure 6, A and B). To evaluate for autoreactive cells undergoing thymic negative selection, we assessed programmed cell death protein 1 (PD-1) on host thymocytes, which has been shown to enable deletion after recognition of self-antigen (40, 41). Indeed, we observed increased PD-1 expression in host thymocytes of prediabetic NOD:B6 mice compared to controls (Figure 5C, Supplemental Figure 6C). While all thymic APCs can mediate negative selection, only mDCs have been shown to enhance production of thymic regulatory T cells (tTregs) in vivo (42). Interestingly, we observed an increase in mDC frequency in NOD:B6 mice that correlated with increased host tTreg frequency, compared to controls (Figure 5D). Consistent with prior reports, enhanced donor tTreg production was not observed in mixed chimeras (14, 18).

**Figure 5.**
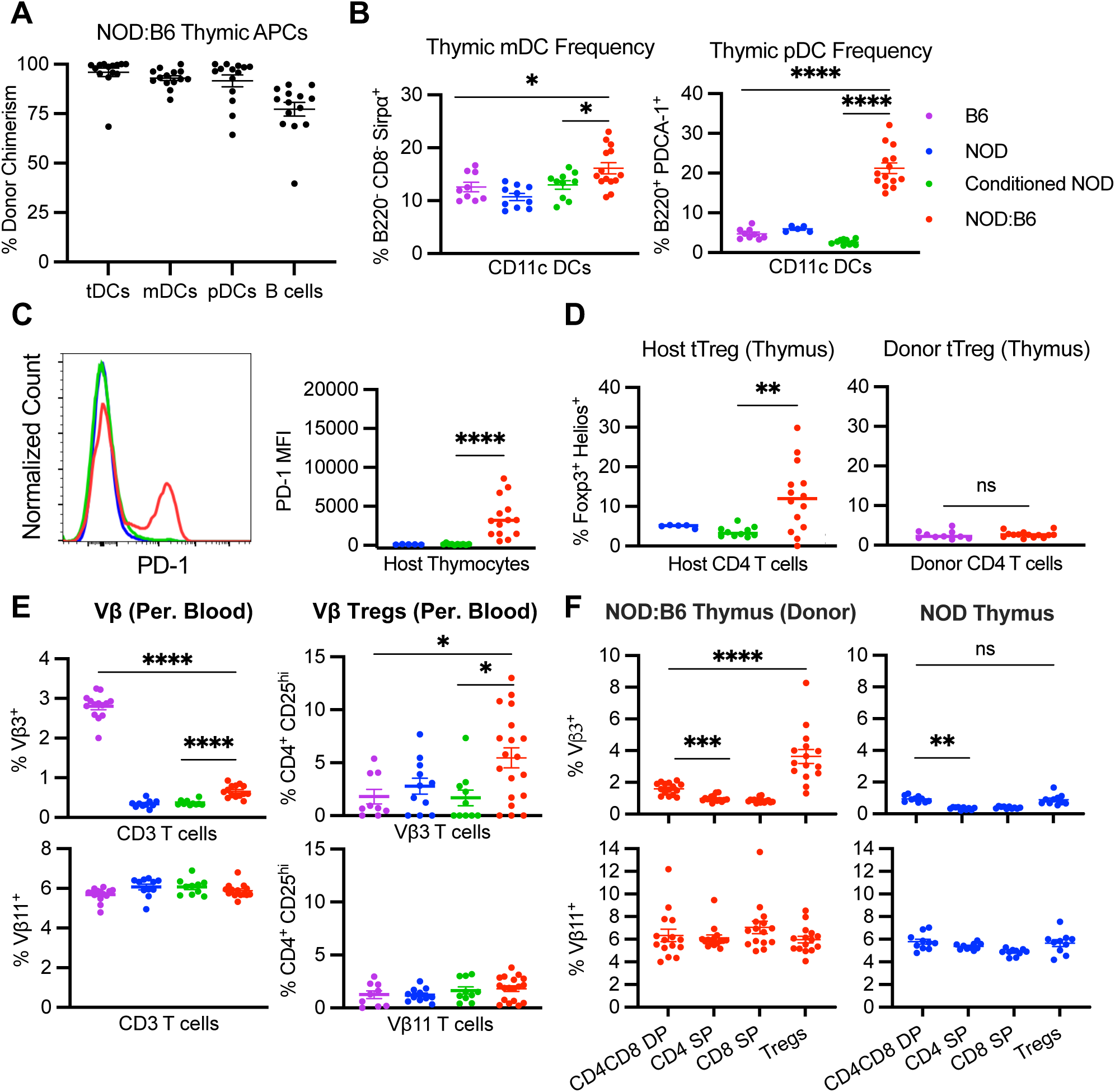
Donor thymic cells are associated with central tolerance and thymic Treg development. **(A)** CD45.2^+^ cell frequency in CD11c^+^ DC subsets including thymus-resident DCs (tDC; CD8^+^ Sirpα^­^), migratory DCs (mDC; CD8^−^ Sirpα^­^), and plasmacytoid DCs (pDC; B220^+^ PDCA-1^+^), as well as B cells 20 weeks post-HCT in prediabetic NOD:B6 mice (*n* = 14). **(B)** mDC and pDC frequency in WT B6, conditioned NOD controls, and prediabetic NOD:B6 (*n* = 9-14). **(C)** Representative histogram and median fluorescence intensity (MFI) of PD-1 expression in CD45.1^+^ thymocytes in prediabetic NOD:B6, NOD, and conditioned NOD control mice (*n* = 9-14). **(D)** CD45.1^+^ thymic Tregs in prediabetic NOD:B6, NOD and conditioned NOD controls (left, *n* = 5-14). CD45.2^+^ thymic Tregs in prediabetic NOD:B6 and WT B6 mice (right, *n* = 9-14). **(E)** Peripheral blood Vβ3^+^ and Vβ11^+^ T cells (left, *n* = 10-15) and Tregs (right, *n* = 10-15) in WT B6, NOD, conditioned NOD controls, and prediabetic NOD:B6 mice (gated on donor cells, red). **(F)** Vβ3 and Vβ11 frequency in thymic T cell subsets of NOD:B6 mice (left, *n* = 15, gated on donor cells) and NOD mice (right, *n* = 10). A-F. Data presented as mean ± SEM. ns = not significant, \**P* < 0.05, \*\**P* < 0.01, \*\*\**P* < 0.001, \*\*\*\**P* < 0.0001. DC = dendritic cell; DP = double positive; SP = single positive.

Although there was no increase in overall donor tTreg production, we still observed tolerance of donor antigens in our mixed chimeras, evidenced by the lack of GVHD. We posited that enhanced donor B6 Treg production would instead be observed in prediabetic NOD:B6 mice specifically for host-reactive T cell clones. To assess this, we measured T cells with the known NOD-reactive Vβ3 domain, which are deleted by genome-encoded superantigen presentation in NOD but not B6 mice (43, 44). Therefore, an untrained B6 T cell repertoire should contain Vβ3^+^ cells, while one trained on NOD antigens should not. As expected, we observed ∼90% deletion of Vβ3^+^ T cells in peripheral blood of NOD and conditioned NOD mice compared to naïve B6 (NOD: 0.34 ± 0.03%, Cond. NOD: 0.37 ± 0.02%, B6: 2.80 ± 0.09%; Figure 5E). In prediabetic NOD:B6 mice, ∼80% of host-reactive donor Vβ3^+^ T cells were depleted, compared to B6, indicating education of donor T cells to host antigens. By contrast, the frequency of Vβ11^+^ T cells, which are not under selective pressure in NOD or B6 strains, was consistent across all controls and mixed chimeras. Although Vβ3 depletion was not complete, we observed a higher frequency of Tregs among Vβ3^+^ donor T cells in mixed chimeras compared to NOD and B6 controls, while Tregs among Vβ11^+^ donor T cells remained consistent (Figure 5E). Vβ3^+^ T cell and Treg frequency differences remained consistent in the peripheral blood of mixed chimeras longitudinally (Supplemental Figure 6D). These observations were further confirmed by analysis of thymic T cell development in prediabetic NOD:B6 mice (Figure 5F). Reduction of donor Vβ3^+^ cell frequency as thymocytes move from double positive CD4^+^CD8^+^ (DP) to single positive (SP) CD4^+^ or CD8^+^ stages can be seen in both mixed chimeras and NOD mice while Vβ11^+^ cells remain consistent throughout. However, thymic Vβ3^+^ donor Tregs are enriched in mixed chimeras but not in NOD mice. Thus, we observed deletion of host-reactive donor (B6) T cells during thymic negative selection and enhanced host-reactive donor tTreg production. In summary, we found evidence for central tolerance mechanisms crucial for GVHD prevention, allograft tolerance, and re-establishment of self-tolerance in mixed chimeric NOD:B6 mice.

### Peripheral tolerance mechanisms are associated with anergy of peripheral host effector cells

Peripheral tolerance mechanisms are essential to suppress allo- and autoreactive T cells that escape thymic selection, and to tolerize remaining conditioning-resistant tissue-resident host T cells. Similar to tTreg frequencies in the thymus after αCD117-based conditioning and HCT, we observed an increase in host tTreg frequencies in the spleens of prediabetic NOD:B6 mice, but no differences in donor tTreg frequencies, as compared to controls (Figure 6A). Furthermore, both host and donor Tregs upregulated the inducible costimulator (ICOS), which is correlated with higher IL-10 secretion and suppressive potential (Figure 6B) (45). Other markers of suppressive activity, CTLA-4 and LAG3 showed no differences (Supplemental Figure 7A). These donor and host tTregs have been shown to mediate peripheral tolerance of allo- and autoreactive T cells through both direct and indirect mechanisms of suppression to promote a tolerogenic state in host dendritic cells (14, 46). In contrast to host conventional dendritic cells (cDCs), we saw an upregulation of the marker programmed death-ligand 1 (PD-L1) on host pDCs (Figure 6C and Supplemental Figure 7B), characteristic of tolerogenic pDCs integral to peripheral tolerance in transplantation and autoimmunity (47, 48). PD-L1/PD-1 signaling between tolerogenic host DCs and host conventional T cells (T_con_) can initiate inhibitory signaling and promote T cell anergy, a crucial tolerance mechanism of autoreactive T cells in the periphery (49, 50). Indeed, we observed an increase in CD73^hi^ FR4^hi^ cells in prediabetic NOD:B6 mice, compared to controls, suggesting anergy in host CD4^+^ T_con_ cells (Figure 6D) (51, 52). To further determine the capacity of these cells to respond to stimulatory signals, we incubated host-type prediabetic NOD:B6 splenic T cells labeled with cell trace violet in vitro with CD3/CD28 stimulation beads at various dilutions (Figure 6E and Supplemental Figure 7C). Although all groups showed a dose-dependent response, cells from conditioned NOD and NOD controls exhibited robust proliferation while mixed chimeric host cells had a blunted response (Figure 6E). Thus, our studies revealed peripheral tolerance mechanisms that promote long-term correction of autoimmunity and allogeneic tolerance in NOD:B6 mice.

**Figure 6.**
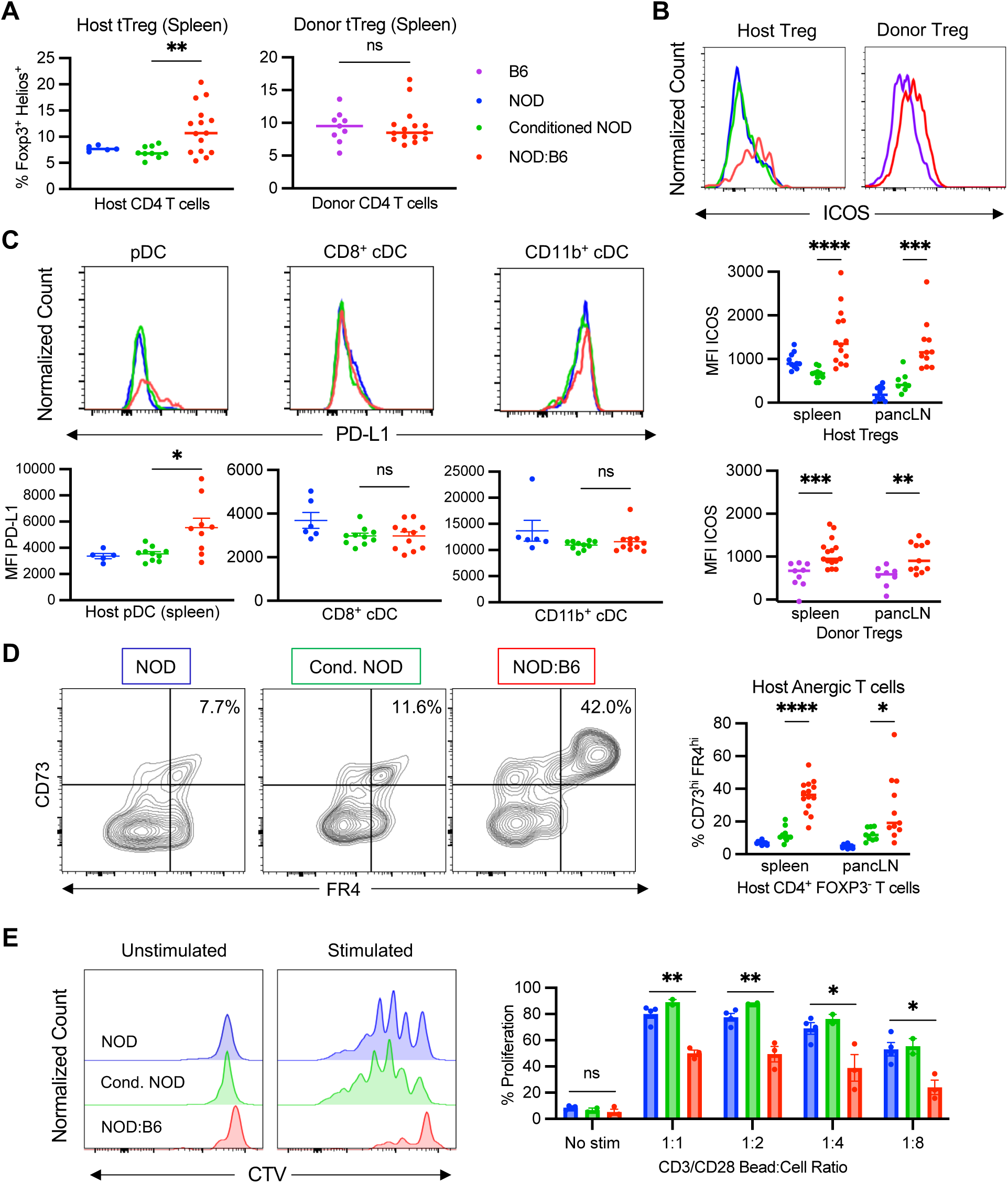
Peripheral tolerance mechanisms are associated with anergy of peripheral host effector cells. **(A)** CD45.1^+^ thymic Tregs (tTregs) in spleens of prediabetic NOD:B6, NOD, and conditioned NOD controls (left, *n* = 5-14). CD45.2^+^ tTregs in spleens of prediabetic NOD:B6 and WT B6 controls (right, *n* = 9-14). **(B)** Representative histogram and median fluorescence intensity (MFI) of ICOS expression in host and donor Tregs in the spleen and pancLN of prediabetic NOD:B6, NOD, conditioned NOD, and WT B6 mice (*n* = 9-14). **(C)** Representative histogram and MFI of PD-L1 of splenic CD45.1^+^ plasmacytoid DCs (pDCs), CD8^+^ conventional DCs (cDCs), and CD11b^+^ cDCs in prediabetic NOD:B6, NOD, and conditioned NOD controls (*n* = 5-10). **(D)** Frequency of CD73^hi^ FR4^hi^ anergic cells among CD45.1^+^ CD4^+^ FOXP3^−^ T_­_cells of prediabetic NOD:B6 spleen and pancLN compared to NOD and conditioned NOD controls (*n* = 9-14). (**E**) Proliferation of host CD4^+^ splenic T cells from prediabetic NOD:B6 compared to NOD and conditioned NOD controls after isolation and incubation in vitro with CD3/CD28 stimulation beads at decreasing dilutions (1:1 to 1:8), assessed with cell trace violet (CTV) dilutions. Statistical analysis done using student’s unpaired t test. A-E. Data presented as the mean ± SEM. ns = not significant, \**P* < 0.05, \*\**P* < 0.01, \*\*\**P* < 0.001, \*\*\*\**P* < 0.0001.

## DISCUSSION

Hyperglycemia and diabetes have been associated with impaired bone marrow engraftment and an altered bone marrow niche (30–33); thus, it was unknown whether mixed chimerism after αCD117-based conditioning was possible in animals with established, fulminant autoimmune diabetes. Here, we reversed autoimmune diabetes in NOD mice using αCD117-based NMA conditioning to achieve durable mixed hematopoietic chimerism and islet tolerance. After diabetes reversal, we observed evidence of central and peripheral tolerance mechanisms resulting in correction of autoimmunity, specific allogeneic donor tolerance, and prevention of GVHD. We also showed that mixed chimerism after αCD117-based conditioning in prediabetic NOD mice eliminated diabetes onset. Thus, we achieved durable mixed chimerism and islet tolerance with this αCD117-based regimen in mice with T1D.

Few studies have described complete reversal of long-standing diabetes in NOD mice (53), and none have led to a clinically-portable approach for established T1D reversal. In one prior study, non-myeloablative conditioning with αCD40L antibody, T cell depletion, 400 cGy TBI, and concurrent transplantation of matched islets, was used to establish mixed hematopoietic chimerism and reverse overt diabetes in NOD mice (15, 16). However, this conditioning regimen relied on tolerization of residual host NOD T cells over a sustained period of weeks, rather than acute bone marrow (BM) elimination and repopulation. Moreover, activated NK/NKT cells present a considerable barrier in co-stimulation blockade-based regimens (54, 55). Additionally, later clinical trials using αCD40L-based conditioning were halted due to thromboembolic complications (56). Zeng and colleagues reported that non-myeloablative conditioning using αCD3 and αCD8 antibodies without radiation supported durable mixed chimerism in diabetic NOD mice (14, 57, 58). However, diabetes reversal relied on host β cell regeneration or preserved islet function in mice with recent-onset diabetes; thus, donor islet tolerance was not addressed in those prior studies. Additionally, the risk of GVHD is high as this approach relies on injection of donor lymphocytes for immunosuppression, known as graft versus host reactivity (11, 59). Protection of remnant islet β cells in humans or NOD mice with recent-onset T1D can be fostered using immune-depleting agents like cyclophosphamide, anti-thymocyte globulin, or autologous HCT (60, 61) resulting in reversal of hyperglycemia, but the general adoption of this approach for T1D, including for chronic T1D, and long-term β cell tolerance remains unestablished. In studies here, mice had diabetes for 6-7 weeks prior to conditioning, HCT and islet transplantation, a duration that precluded significant host β cell preservation (Figure 3, E and I).

Human αCD117 (briquilimab) has shown promising results without systemic toxicity in clinical trials for several stem cell diseases (62–66). As in our prior work with diabetic B6 mice (18), αCD117 achieved targeted bone marrow niche clearance without systemic toxicity in NOD mice. However, unlike our prior work in B6 mice, a regimen combining αCD117 with 225 cGy TBI and anti-CD4/8 T cell depletion was unsuccessful in establishing mixed chimerism with NOD mice (Supplemental Figure 1B). To build our NMA regimen, we systematically evaluated conditioning-resistant cell populations and considered possible autoimmune barriers to engraftment. Although JAK inhibition alone did not provide sufficient immune suppression, baricitinib alongside monoclonal antibodies create a favorable environment for engraftment in the setting of autoimmunity (Supplemental Figure 1C). JAK inhibitors were shown to enhance HSPC engraftment and allotolerance in mice and non-human primates (22, 27), are in clinical trials for acute and chronic GVHD, and are FDA-approved to treat autoimmune rheumatoid arthritis (67). In preclinical studies, JAK1/2 inhibitors were shown to alleviate the autoreactive T cell burden and inflammation of T1D (24), and in clinical trials delayed progression in new-onset T1D (68). However, incorporation of JAK inhibitor in NMA conditioning without HCT did not reliably prevent diabetes in NOD mice (Figure 2A). To prevent or correct diabetes in NOD mice with HCT, we speculate that baricitinib is useful for NMA conditioning to promote allogeneic immune suppression, and also for overcoming an autoimmune engraftment barrier in T1D.

GVHD risk is a challenge during HCT. Lineage-depleted HSPCs can reduce GVHD risk (69) and can achieve mixed chimerism with our conditioning (Supplemental Figure 1D). However, unmanipulated bone marrow or peripheral mobilized stem cell transplantation is the current clinical standard. We did not observe evidence of GVHD in NOD:B6 chimeras transplanted with unmanipulated whole bone marrow (WBM), showing that use of T cell depleting antibodies and JAK1/2 was sufficient in the setting of a replete graft (67). Moreover, successful use of either HSPCs or WBM to achieve mixed chimerism consistently indicates that our conditioning regimen does not rely on donor T cells for HSC engraftment.

Conditioning radiation needed to achieve mixed chimerism can be associated with significant toxicity in pre-clinical or clinical settings (70). Prior studies reported dosages of 400-1100 cGy TBI for conditioning diabetic NOD mice to achieve mixed chimerism (12, 13, 15, 16). 400 cGy (LD_50_) is a recognized dosage associated with radiation side effects, including bleeding, renal medullary damage, and risk of secondary malignancy (71, 72). In contrast, 225-250 cGy is associated with transient signs of hematopoietic impairment and lower risk of secondary malignancy. Here, we report successful mixed NOD:B6 chimerism following TBI reduction to 225-250 cGy combined with αCD117 conditioning, transient T cell depletion and JAK1/2 inhibition. Moreover, mice treated solely with this NMA conditioning recovered hematopoiesis without any transplant, transfusion, additional supportive care, diet supplement, or other interventions (Supplemental Figure 2C). Reduction of TBI to <225 cGy did not consistently promote NOD:B6 hematopoietic chimerism (Supplemental Figure 1E), establishing a lower limit on TBI dose in this NMA conditioning regimen. Diabetic NOD mice have inflammatory and microvascular changes in the bone marrow niche (30, 32, 73), and prior reports have established the difficulty of achieving BM engraftment and eradication of host lymphocytes in NOD mice, especially with NMA regimens using reduced TBI doses (25). We speculate that modification of NMA conditioning in future studies, like TBI dosage fractionation, TLI, prolonged JAK inhibition, or addition of other conditioning reagents to target inflammatory factors in the bone marrow niche (11, 22, 74, 75) could further reduce or eliminate radiation toxicity during NOD mouse conditioning.

To assess mechanisms of allotolerance and autoimmunity suppression in NOD mice with mixed chimerism, we characterized cellular and molecular features of central and peripheral tolerance. We observed consistent overall and cell type-specific mixed chimerism in multiple compartments, including bone marrow, spleen, thymus and circulating blood. Additionally, the data support the view that thymic-mediated central tolerance, including evidence of donor APC chimerism (DC subsets and B cells) and elimination of host-reactive Vβ3^+^ donor T cells in the thymus, or conversion of these cells to Tregs, promote allogeneic tolerance in these settings. Tetramer analysis revealed a reduction or elimination of peripheral autoreactive CD8 cells. Lack of islet immune cell infiltration in mixed chimeric NOD mice after HCT and islet transplantation, suggests that donor alloantigen tolerance and reversal of autoimmunity was achieved using mixed chimerism. Likewise, mixed chimerism prevented T1D onset in prediabetic NOD mice, and adoptive transfer studies with these mice confirmed durable correction of autoimmunity. Maintenance of immune competence after NMA conditioning and islet transplantation to achieve diabetes reversal is an important goal for clinical translation. We observed brisk recovery of blood counts after HCT and robust rejection of third-party FVB islet grafts by mixed chimeric NOD:B6 mice, evidence of immune function.

These studies in inbred rodent models are potentially limited by the possibility of heterologous immune responses (76), which could impact immunological tolerance, bone marrow engraftment, and autoreactivity. Additionally, although rejection of third-party grafts can assess immune reconstitution, future studies could evaluate additional immune responses to non-MHC expressing pathogens or the heterogeneity of the reconstituted repertoire. NOD mice are an important model of T1D, but have known limitations (77), including sexually dimorphic diabetes risk, and studies here are limited by a focus on NOD female mice. Larger animal studies, such as those that established the use of αCD117 for bone marrow conditioning prior to clinical trial, may be necessary to evaluate conditioning kinetics of bone marrow niche clearance, long-term allotolerance, and correction of autoimmunity. Despite challenges of obtaining donor-matched solid organs and bone marrow, success with combined renal and bone marrow transplantation (78) supports the feasibility of this approach. We show that islets could be transplanted months after HCT, suggesting that future studies could establish the scheduling flexibility of conditioning, islet transplantation and HCT steps in our regimen, using bone marrow and islets isolated from a single donor, including the possibility of cryopreserved islets (79).

In summary, this work identifies a novel AMA conditioning regimen for bone marrow engraftment in a model of autoimmune diabetes. A sequence of αCD117 antibody and baricitinib-based NMA conditioning followed by allogeneic WBM and islet transplantation achieved durable hematopoietic chimerism, islet allograft tolerance, suppression of islet autoimmunity, and reversed type 1 diabetes, without GVHD. In principle, these findings could apply to tolerance induction of other tissues and cells, including transplanted stem cell-derived replacement β cells.

## METHODS

### Sex as a Biological Variable

Our study examined female mice due to the very low incidence of autoimmunity (< 5%) in the male population in our colony, in line with prior reports. Our prior studies of mixed chimerism and diabetes do not show sex-dimorphic effects (18).

### Study Design

The objective of this research was to expand a low-toxicity conditioning and transplantation approach from prior mouse models into an autoimmune setting, with the hypothesis that the approach will allow for simultaneous correction of autoimmunity, solid-organ transplant tolerance, and diabetes reversal. NOD mice were chosen as a model of autoimmunity due to the extensive similarities in pathogenesis to human T1D which allow for study of mismatched bone marrow transplantation. Animals were randomly assigned to experimental groups, and all samples represent biological replicates. The primary endpoint of the controlled laboratory study was 20 weeks after HCT, loss of islet graft tolerance, or severe GVHD, whichever presented first. Serial immune monitoring was performed on peripheral blood every 4 weeks to assess mixed chimerism levels, autoreactivity, and host/donor tolerance, confirmed by endpoint splenic, thymic, bone marrow, lymph node, and peripheral blood analysis. Weekly blood glucose monitoring was performed to evaluate for diabetes from graft loss or autoimmunity, confirmed by endpoint islet histology. Weekly monitoring of weight and skin changes were used to evaluate occurrence of GVHD, confirmed by endpoint intestinal histology. Initial engraftment success was defined as at least 5% chimerism in both myeloid and lymphoid lineages in whole blood, and endpoint engraftment success defined by at least 5% chimerism in LSK HSCs of the bone marrow.

### Animals

Female NOD (7-8 weeks old; Stock #:001976) and NOD Rag1KO (6-7 weeks old; Stock #:003729) were used as recipients in transplantation and adoptive transfer studies, respectively. Female B6 (Stock #:000664), FVB (Stock#:001800), or B6 FOXP3 DTR (Stock#:016958) donor mice were used for bone marrow (6-7 weeks old) and islet (10-16 weeks old) isolations. All mice were purchased from The Jackson Laboratory, fed standard chow and water ad libitum, and housed in specific-pathogen-free barrier facility at the Stanford School of Medicine. Animals receiving transplantation were fed trimethoprim and sulfadiazine (Uniprim) anti-bacterial diet for a period of 6 weeks, starting two weeks prior to transplantation conditioning.

### Diabetes monitoring and maintenance

All animals were monitored weekly for weight and blood glucose using True Metrix Blood Glucose Monitor and Test Strips (Trividia Health) and were considered diabetic after two consecutive readings above 250 mg/dL. Mice are considered euglycemic when their non-fasting blood glucose returns to <250mg/dL. Diabetic animals that were maintained for further experiments or before transplantation were given insulin implants as needed (Linshin), dosed by weight and inserted according to manufacturer’s instructions. The insulin release is approximately 0.1U/24hr/implant for 30 days. Diabetic mice may be temporarily treated with insulin implants and/or supplemental insulin glargine (Sanofi) in the week prior to and after transplantation to achieve normoglycemia.

### Conditioning, reagents, and equipment

A graphical timeline of conditioning is shown in Figure 1B and 3A. Mice were given 500μg diphenhydramine HCl i.p. approximately 10-15 minutes prior to αCD117. 400-500μg αCD117 was injected retro-orbitally into mice under isoflurane anesthesia on day −4 prior to HCT. Mice were irradiated on day −2 with 225-250 cGy TBI. 600μg of αCD4 and 300μg αCD8 was administered i.p. on days −2 through 2 and day 7. 400μg JAK1/2 inhibitor baricitinib was administered s.c. days −3 through 8, prepared as described previously (22). ACK2 αCD117 mAb was purchased from BioLegend αCD4 (GK1.5) and αCD8 (YTS169.4) were purchased from Bio X Cell. Diphenhydramine HCl and baricitinib were purchased from MedChem Express. Animal irradiation (XRT) was performed using a Kimtron Polaris IC-250 Biological Irradiator with a 225 kV X-ray tube filtered by 0.5mm Cu source set at 225kV, 13.3mA. Mice were divided in irradiation pie cages from Braintree Scientific when irradiated.

### CBC

Complete blood counts with differential were performed by Stanford Veterinary Service Center Diagnostic Laboratory using standard equipment. Automated hematology was performed on the Sysmex XN-1000V hematology analyzer system. Blood smears were made for all CBC samples and reviewed by a clinical laboratory scientist. Manual differentials were performed as indicated by species and automated analysis. Reference ranges were as follows: WBCs, 1.8–10.7 K/μL; RBCs, 6.36-9.42 M/μL; hemoglobin 13.7-16.4 g/dL, hematocrit, 39%–47%; neutrophils, 0.1-2.4 K/μL; lymphocytes, 0.9-9.3 K/μL. CBCs from female untreated B6 (*n* = 9) and prediabetic or diabetic NOD mice (*n* = 5 each) mice were used for the strain-specific averages.

### Bone marrow isolation, enrichment, and transplant

Sex-matched donor B6 or FVB mice or (6-7 weeks old) were euthanized and femurs, tibias, and vertebral bodies were collected. Bones were crushed via mortar and pestle in PBS with 2% FBS and 10mM HEPES to recover WBM. WBM was filtered through a 70-µm cell strainer, and RBCs were lysed in RBC Lysis Buffer (BioLegend). WBM cells were stained with Trypan Blue (StemCell Technologies) and counted with a Countess 3 Automated cell counter (ThermoFisher Scientific). Lineage-negative (Lin^−^) bone marrow cells were enriched by magnetic column separation using a Lineage Cell Depletion cocktail (Miltenyi Biotec) as per manufacturer’s instructions. 2.5E6 Lin^−^ cKit^+^ cells or 30E6 WBM cells suspended in 100 µl of PBS were injected retro-orbitally. Lin^−^ cKit^+^ HSPC preparation composition has been previously described (18).

### Islet isolation and transplantation

Islet isolation and transplantation was performed as previously described with minor modifications (80, 81). Briefly, pancreases are perfused with 100-125μg/mL Liberase TL (Roche Diagnostics) through the common bile duct and digested in a 37°C water bath for 18-22 minutes. After washing with Hank’s Buffered Saline (HBS; Caisson Labs), the crude digest is purified over a discontinuous density gradient, washed once more with HBS, and cultured overnight in 5.5 mM glucose RPMI 1640 (Corning) supplemented with 10% FBS, 10mM HEPES, and 1% penicillin-streptomycin solution. Recipient mice were anesthetized with a ketamine/xylazine mix and given subcutaneous analgesics. 200-400 islets were injected under the kidney capsule of recipient mice with a micro-capillary, as described (82). The nephrectomy procedure involved the same anesthetic regimen as islet transplantation and renal vessels were first tied to prevent hemorrhage before the kidney containing islet graft was removed.

### Histology

Islet graft-bearing kidneys, pancreases, and intestines were fixed in 4% paraformaldehyde overnight, incubated overnight in 30% sucrose, embedded in optimal cutting temperature compound, and frozen on dry ice. 6-10μm sections were made on a Leica CM3050 S (Leica Biosystems). Immunofluorescent staining was performed using standard methods. Briefly, sections were blocked for 1hr then incubated with primary antibodies overnight at 4°C. Sections were washed for 5 minutes x 3 before incubation with secondary antibodies for 2 hours at room temperature or overnight at 4°C and washed 3 x 5 minutes again. Slide covers were set with Hard-set Mounting Medium with DAPI (Santa Cruz Biotechnology). Slides were imaged on Zeiss AxioM1 or Leica SP2 confocal microscopes. Post-processing and color channel merging was performed in Fiji (http://fiji.sc/) (83). Primary and secondary antibodies and dilutions are documented (Supplemental Table 1). For histological examination of adoptive transfer recipients, pancreases were removed, fixed in 10% buffered formalin, embedded in paraffin, sectioned, and stained with hematoxylin and eosin by Histo-Tec Laboratory.

### Peripheral blood, spleen, thymus, pancLN, and BM preparation for flow cytometry

100 µl of whole blood was collected from the facial vein using Goldenrod 5mm animal lancet into EDTA coated tubes (BD). Spleens, thymuses, and pancreatic lymph nodes were directly homogenized through a 70µm cell strainer with a syringe plunger. BM cells were isolated as above. Samples underwent RBC lysis in RBC Lysis Buffer (BioLegend) for 5-10 min at 4°C before downstream staining for analysis.

### Flow cytometry analysis

Gating strategies can be found in figure legends, Supplemental data (Supplemental Figure 8), or described previously (18). For analysis of mixed chimerism, cells were first stained with LIVE/DEAD Fixable Near-IR Dead Cell Stain Kit (ThermoFisher Scientific) and blocked with TruStain FcX anti-mouse (Biolegend) for 10 min on ice in Cell Stain Buffer (Biolegend). Extracellular markers were stained with antibodies listed (Supplemental Table 2) at manufacturer’s recommended dilutions. Tetramers (NIH Tetramer Core Facility) were stained with extracellular marker antibodies (0.014 mg/mL). Peptide sequence for IGRP is: VYLKTNVFL_206-214_. An irrelevant RSV control tetramer (KYKNAVTEL, H-2K^d^) was used to validate the specificity of the IGRP tetramer staining (Supplemental Figure 5A). Staining of intracellular markers (Supplemental Table 2) was conducted with Biolegend True-Nuclear Transcription Factor Buffer Set as per manufacturer’s instructions. Cells were analyzed with a 5L Aurora (Cytek Biosciences) and data were analyzed using FlowJo (10.10).

### Adoptive transfer

NOD Rag1KO female mice at 6-7 weeks old were used as diabetes-susceptible recipients. Donor mice were naïve prediabetic 12-week-old female NOD, diabetic NOD between 14-18 weeks of age, prediabetic mixed chimeras at 12 weeks post-HCT or diabetic mixed chimeras at 20 weeks post-HCT. Splenocytes were isolated from donor mice as described above and sorted using a MojoSort Mouse CD3 T cell Isolation Kit and MojoSort Magnet (Biolegend). Experiments with diabetic donors used 10E6 whole splenocytes per recipient while experiments with prediabetic mice used 2E6 magnetically sorted CD3^+^ cells per recipient. Cells were resuspended in 100ul of PBS injected retro-orbitally in recipients under isoflurane anesthesia.

### In vitro proliferation assay

Cells were isolated from the spleen of prediabetic NOD at 12-14 weeks old, conditioned prediabetic NOD 12-14 weeks post-conditioning, and NOD:B6 12-14 weeks post-HCT as described in splenic preparation for flow cytometry. T cells were separated using EasySep Mouse T cell Isolation Kit (STEMCELL technologies) and labeled with CellTrace Violet dye (Thermo Fisher Scientific) in PBS for 20 min at 37C per manufacturer instructions. Next, T cells (100k) were incubated per well of a 96-well plate with CD3/CD28 stimulation beads at serial dilutions (1:1 to 1:64) for 5 days. Cells were collected, stained, and analyzed by flow cytometry as described above.

### Statistical analysis

Statistical details of all experiments can be found in the figure legends and results section, including *n* values. All data are presented as means ± SEM, where *n* represents number of animals, unless otherwise noted. Mice with accidental mortality due to handling were excluded from analysis. Statistical analysis was performed using Prism 10 (GraphPad). Differences between the means of two groups with *n* < 20 were tested with Mann-Whitney U test for unpaired samples or Wilcoxon matched-pairs sign rank test for paired samples, unless otherwise stated. Kaplan-Meier survival curve comparisons were tested with log-rank Mantel-Cox as well as Fisher’s exact tests where noted. For contingency tables with sample size *n* > 40, χ^2^ test was used. A *P* value of < 0.05 was considered statistically significant. * *P* < 0.05, \*\**P* < 0.01 *** *P* < 0.001, \*\*\*\**P* < 0.0001

### Study Approval

Animal experiments were approved by the Stanford Administrative Panel on Laboratory Animal Care, in line with ARRIVE guidelines.

## Data Availability

All data generated or analyzed during this study are included in this published article (and its supplemental information files) or are available from the corresponding author on reasonable request. The supporting data values file includes values underlying graphed data and reported means presented in both the main text and supplemental material.

## Author contributions

P.B. contributed to conceptualization, designed and performed experiments, data collection, data analysis and visualization, wrote the manuscript, and was involved in funding acquisition. S.A.R. advised on experimental design and performed experiments, data collection, data analysis and visualization, edited the manuscript, and was involved in funding acquisition. B.R. performed experiments, data collection, data analysis, and visualization, and was involved in funding acquisition. Y.M. and Y.Z. performed experiments, data collection, and data analysis. J.P., B.J.V., S.P., C.A.C, and H.K. advised on experimental design and performed experiments, data collection, and data analysis; X.G., N.N, W.Z., R.R., D.B., and M.A.M performed experiments and data collection; E.M. and J.A.S. provided guidance and feedback on experimental design, results, provided reagents, reviewed and edited the manuscript, and were involved in funding acquisition. S.K.K. contributed to conceptualization and experimental design, assisted with data analysis and visualization, wrote the manuscript, supervised the project, acquired funding, and is the guarantor of this work.

## Acknowledgments

We thank members of the Kim and Meyer groups, especially Drs. Y. Hang, R. Whitener, S. Park, B. Iliopoulou, and C. Bader for experimental advice and encouragement. Flow cytometry analysis was done in the Stanford Shared FACS Facility, with special thanks to core director, Dr. Lisa Nichols. CBC testing was done by the Stanford Animal Diagnostic Laboratory. We thank the Stanford Diabetes Research Center (SDRC) Islet Core for assistance with islet transplantation. P.B. is a student in the Medical Scientist Training Program (MSTP) and PhD Program in Immunology at Stanford. P.B. is supported by the NIH (T32 GM736543) and Stanford Interdisciplinary Graduate Fellowship through Bio-X (Morgridge Family Fellow). S.A.R. is supported by Institute for Immunity, Transplantation and Infection – Stanford Autoimmunity & Allergy Supergroup and NIH grant LAUNCH 1TL1DK139565-0. B.R. is supported by the VPUE Research Fellowship at Stanford. Work in the Kim group was supported by the Breakthrough T1D Northern California Center of Excellence (S.K.K, J.A.S, E.M), NIH awards (R01 DK107507; R01 DK108817; U01 DK123743; P30 DK116074 to S.K.K), the Reid Family, H.L. Snyder Foundation and Elser Trust, and the Stanford Diabetes Research Center (SDRC). Work here was also supported by the Islet Research Core in the SDRC.

## Notes

**Conflict of Interest:** JAS is a co-founder, stockholder, and board member, HSK is an employee and stockholder, and CAC is a stockholder of Jasper Therapeutics, Inc. S.A.R is a consultant and stockholder of Tolerance Bio, Inc.

### Competing Interest Statement

JAS is a co-founder, stockholder, and board member, HSK is an employee and stockholder, and CAC is a stockholder of Jasper Therapeutics, Inc. S.A.R is a consultant and stockholder of Tolerance Bio, Inc.

