## Supplemental Materials for "Curing autoimmune diabetes with islet and hematopoietic cell transplantation after CD117 antibody-based conditioning"

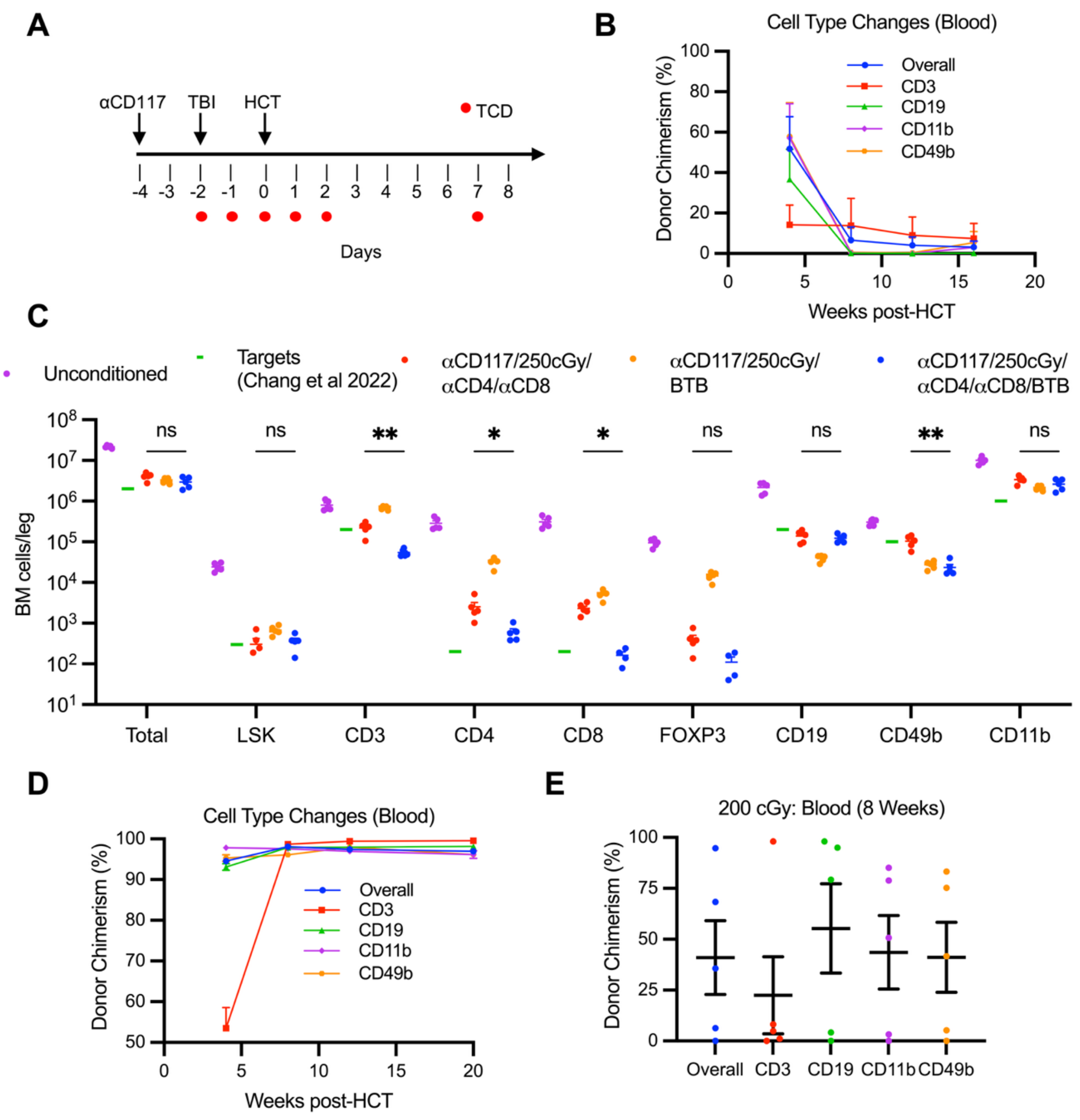


**Supplemental Figure 1. Non-myeloablative conditioning requires JAK inhibition to permit durable allogeneic donor chimerism in a radioresistant NOD model. (A)** Reduced intensity conditioning regimen without JAK inhibition. 500µg αCD117 is administered on day -4, 300cGy TBI on day -2, 600µg/300µg of αCD4/CD8 on days -2 through +2 and day +7. On day 0, mice are transplanted with 30E6 WBM cells or 1.5E6 HSPCs after TCD treatment. Mixed chimerism is analyzed every 4 weeks afterwards, in peripheral blood. (**B**) Longitudinal chimerism analysis of peripheral blood after conditioning (without JAK inhibition) and HCT depicting overall, CD3$^+$ T cell, CD19$^+$ B cell, CD11b$^+$ myeloid cell, and CD49b$^+$ NK cell donor (CD45.2$^+$) chimerism ($n$ = 5). (**C**) Bone marrow (BM) absolute cell count analysis at Day 0 after conditioning regimen containing αCD117, 250 cGy TBI, and αCD4/CD8 with (blue, Figure 1A) or without (red, Supplemental Figure 1A) JAK inhibitor baricitinib (BTB), as well as αCD117, 250 cGy TBI, and BTB without TCD (orange). Approximate target values for depletion are shown in green from studies in C57BL/6 mice in Chang et al 2022, and unconditioned NOD in purple. (**D**) Chimerism analysis of peripheral blood over 20 weeks post-HCT with 2.5E6 B6 HSPCs and conditioning with JAK inhibition ($n$ = 5). (**E**) Chimerism analysis of peripheral blood at 8 weeks post-HCT with 30E6 B6 WBM cells and conditioning with JAK inhibition and XRT reduced to 200 cGy ($n$ = 5). Data are represented with mean ± SEM. TBI = total body irradiation; HCT = hematopoietic cell transplant; TCD = T cell depletion; HSC = hematopoietic stem cell; LSK = Lin$^-$ cKit$^+$ Sca-1$^+$

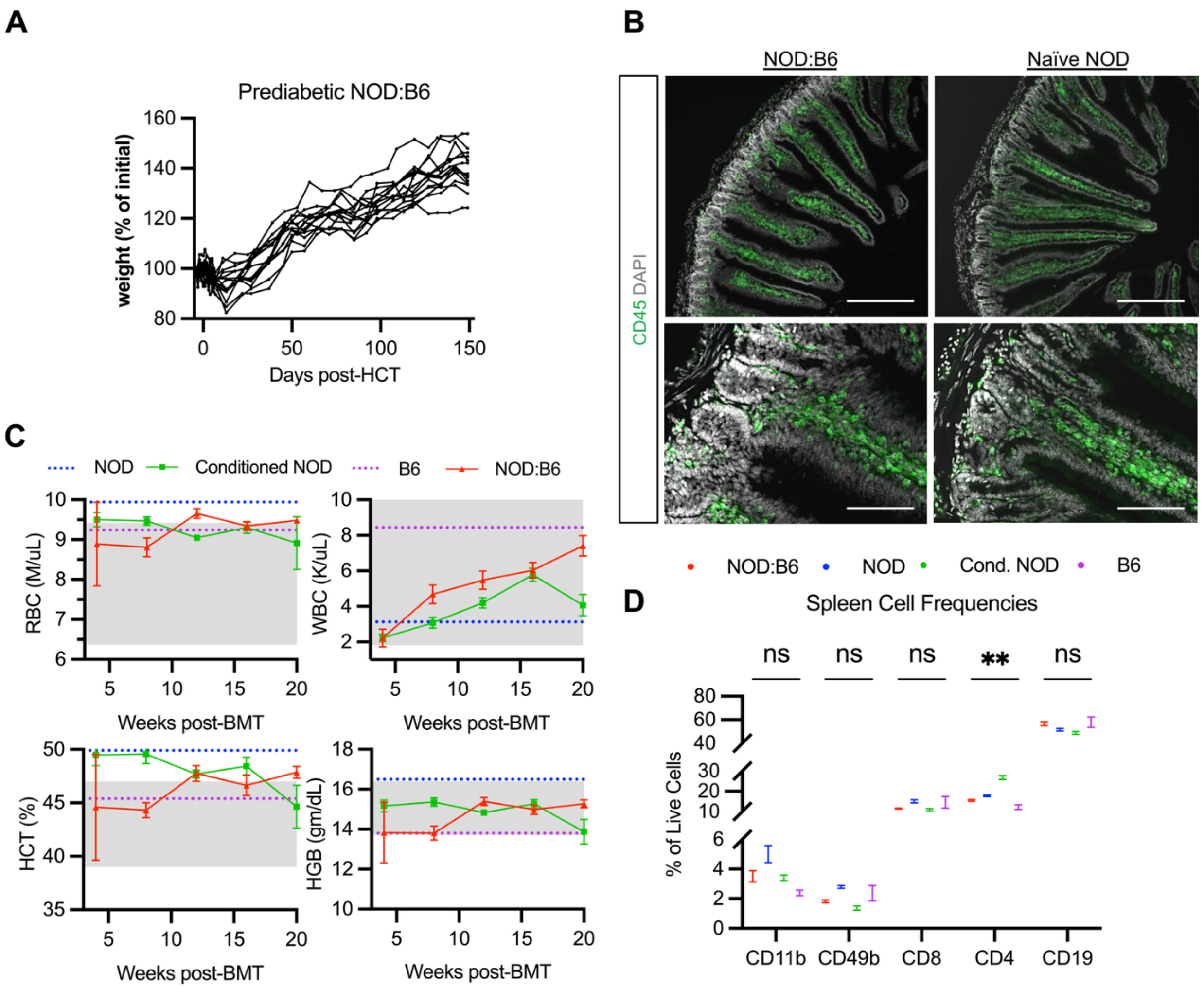


**Supplemental Figure 2. Functional status of prediabetic mixed chimeras after non-myeloablative conditioning and HCT.** (**A**) Weight after HCT of prediabetic NOD:B6 as a percentage of initial weight prior to conditioning start (*n* = 15, sum of 3 independent experiments). (**B**) Representative host duodenal histology of prediabetic NOD:B6 chimeras (*n* = 14) at 20 weeks post-HCT compared to naïve NOD, stained for CD45. Scale bars = 200µm (top) or 50µm (bottom). (**C**) Complete blood count values for prediabetic NOD:B6 mice after conditioning and HCT (red; n = 10) vs. prediabetic NOD mice after conditioning only (green; n = 10). Blue dotted lines show averages from naïve prediabetic NOD mice (n = 5) and purple dotted lines show averages from naïve B6 mice (n = 9). Gray boxes show reference ranges as provided with hematology analyzer equipment (see Methods). WBC = white blood cell; RBC = red blood cell; HGB = hemoglobin; HCT = hematocrit. (*n* = 5-10). (**D**) Myeloid, NK, CD8+ T, CD4+ T, and B cell frequencies in spleens of prediabetic NOD:B6, naïve NOD, conditioned NOD, and naïve B6 mice (*n* = 9-14). Data are represented with mean ± SEM. *P* values were calculated using Mann-Whitney tests. $^{**}P < 0.01$; ns = not significant

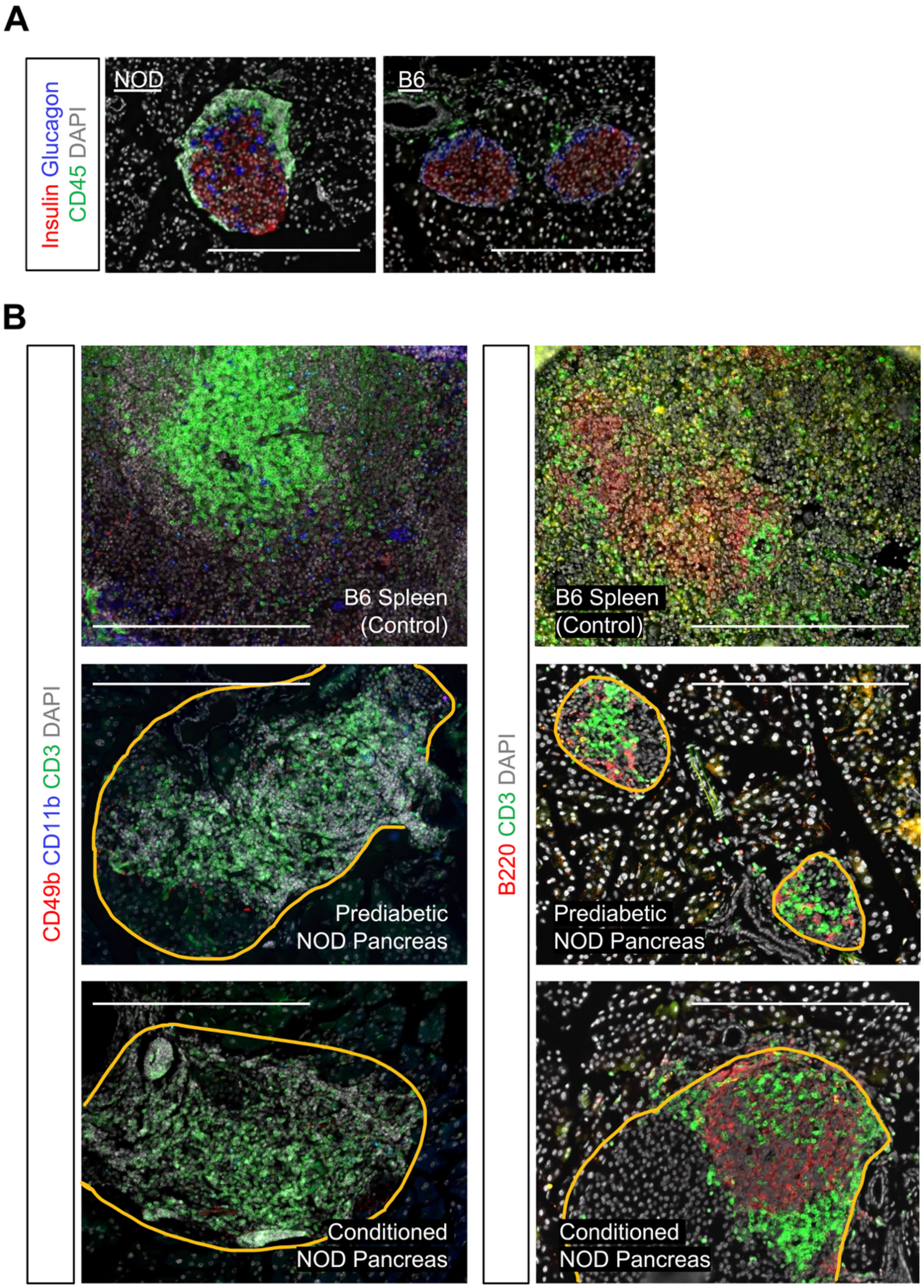


**Supplemental Figure 3. Insulitis is present in conditioned mice but not in mixed chimeras.** (**A**) Representative pancreatic histology of prediabetic NOD at 8 weeks compared to naïve B6 after immunostaining for insulin, glucagon, and CD45. ($n$ = 4-6; scale bars = 200μm). (**B**) Representative pancreas histology of naïve prediabetic NOD at 12 weeks, or conditioned prediabetic NOD (20 weeks after conditioning), immunostained as indicated for CD3, CD49b and CD11b, or CD3 and B220 ($n$ = 3 each, scale bars = 200μm, islets circled in yellow). B6 spleen included as a staining control.

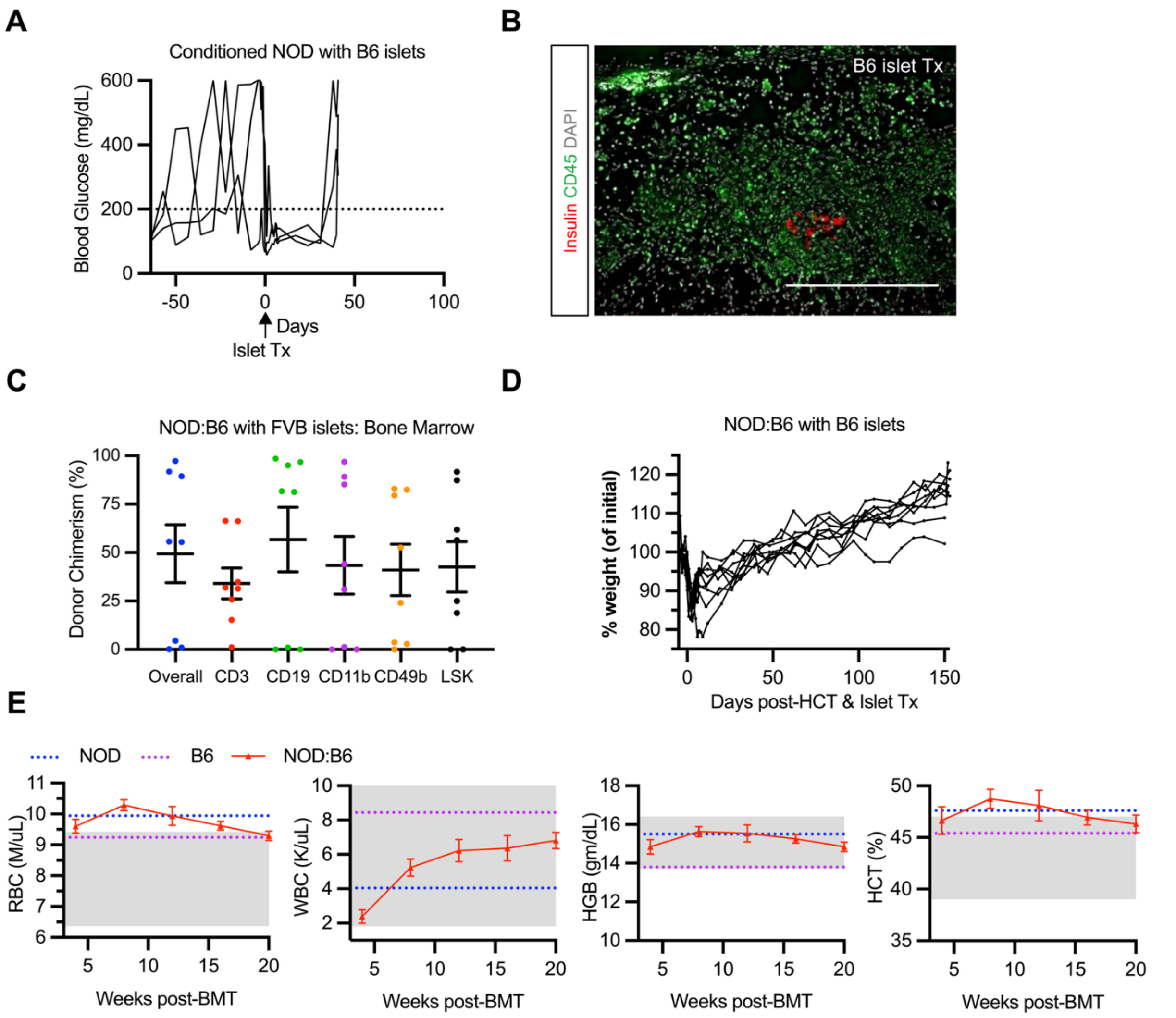


**Supplemental Figure 4. Functional status of diabetic mixed chimeras after non-myeloablative conditioning, HCT, and islet transplantation. (A)** Non-fasting blood glucose of conditioned diabetic NOD mice without HCT that received B6 islets (n = 3). Dotted line (200 mg/dL) indicates normoglycemia threshold. **(B)** B6 islet graft 5 weeks after transplantation in conditioned diabetic NOD, stained for insulin and CD45 (n = 3). Scale bar = 200µm (**C**) Bone marrow chimerism at endpoint (islet rejection) in diabetic NOD:B6 mice that received FVB islets, including $Lin^-$ $cKit^+$ $Sca\text{-}1^+$ (LSK) HSC chimerism (*n* = 8, sum of 3 independent experiments). (**D**) Weight after HCT and B6 islet transplantation of diabetic NOD:B6 as a percentage of initial weight prior to conditioning start (*n* = 11, sum of 3 independent experiments). (**E**) Complete blood count values for diabetic NOD:B6 mice after conditioning, HCT, and B6 islet transplantation (red, *n* = 9). Blue dotted lines show averages from naïve diabetic NOD mice (*n* = 5) and purple dotted lines show averages from naïve B6 mice (*n* = 9). Gray boxes show reference ranges as provided with hematology analyzer equipment (see Methods). WBC = white blood cell; RBC = red blood cell; HGB = hemoglobin; HCT = hematocrit. Data are represented with mean ± SEM.

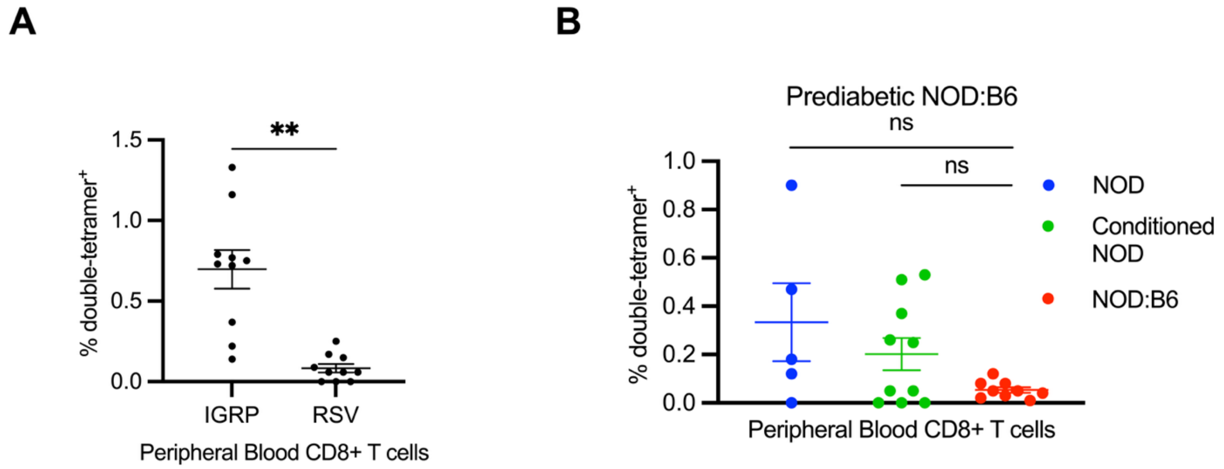


**Supplemental Figure 5. Autoreactive T cells can be tracked using islet autoantigen specific tetramer analysis.** (**A**) Islet autoantigen IGRP-double-tetramer$^+$ frequency compared to irrelevant RSV-double-tetramer$^+$ frequency among peripheral blood CD8$^+$ T cells in diabetic NOD ($n$ = 10). (**B**) Frequency of IGRP-double-tetramer$^+$ autoreactive cells among CD3$^+$ CD8$^+$ T cells in peripheral blood of prediabetic NOD and prediabetic NOD:B6 at 20 weeks post-conditioning and HCT ($n$ = 5-10). Data are represented with mean ± SEM. P values were calculated using Wilcoxon matched-pairs signed rank test (A) or Mann-Whitney tests (B). **$P$ < 0.01; ns = not significant

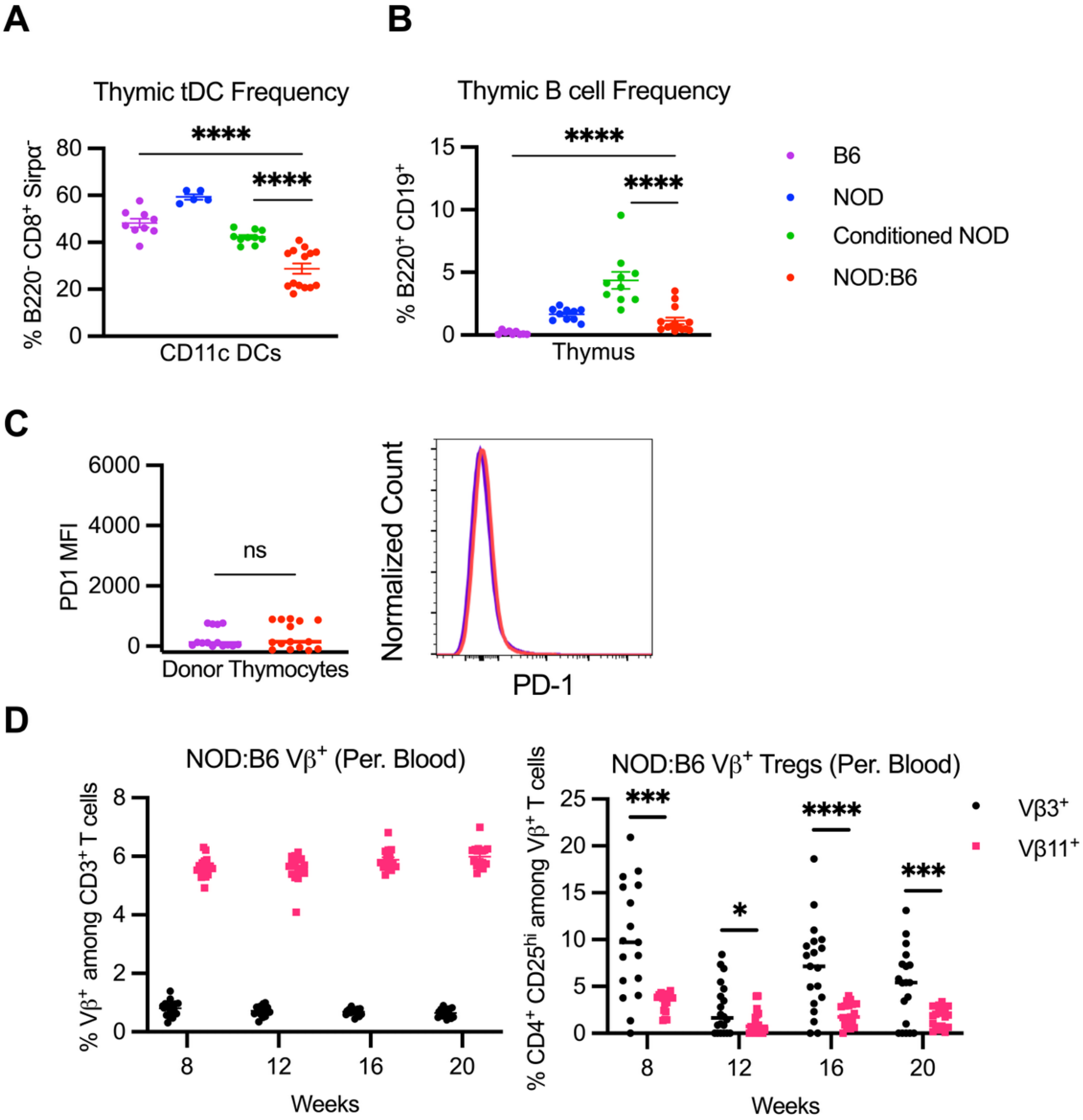


**Supplemental Figure 6. Donor antigen presenting cell presence is associated with thymic deletion of host and donor T cells.** (**A**) Frequency of thymus-resident DCs (tDC; B220$^-$ CD8$^+$ Sirpα$^-$) among CD11c$^+$ DCs in naïve B6 controls, naïve prediabetic NOD, conditioned prediabetic NOD controls at 20 weeks post-conditioning, and prediabetic NOD:B6 at 20 weeks post-conditioning and HCT (*n* = 9-14). (**B**) Frequency of B cells among thymocytes in naïve B6 controls, naïve prediabetic NOD, conditioned prediabetic NOD controls at 20 weeks post-conditioning, and prediabetic NOD:B6 at 20 weeks post-conditioning and HCT (*n* = 9-14). (**C**) Representative histogram and mean ± SEM of median fluorescence intensity (MFI) of PD-1 expressed by CD45.2$^+$ donor thymocytes in prediabetic NOD:B6 compared to B6 controls (*n* = 9-14). (**D**) Frequency of Vβ3 and Vβ11 among CD3$^+$ T cells in peripheral blood of prediabetic NOD:B6 from 8 to 20 weeks post-HCT (left). Frequency of CD4$^+$ CD25$^{hi}$ Tregs among Vβ3$^+$ or Vβ11$^+$ T cells in peripheral blood of prediabetic NOD:B6 from 8 to 20 weeks post-HCT (right) (*n* = 14). Data are represented with mean ± SEM. P values were calculated using Mann-Whitney tests (A-C) or Wilcoxon matched-pairs signed rank test (D) *$P < 0.05$, **$P < 0.01$, ***$P < 0.001$, ****$P < 0.0001$. DC = dendritic cell

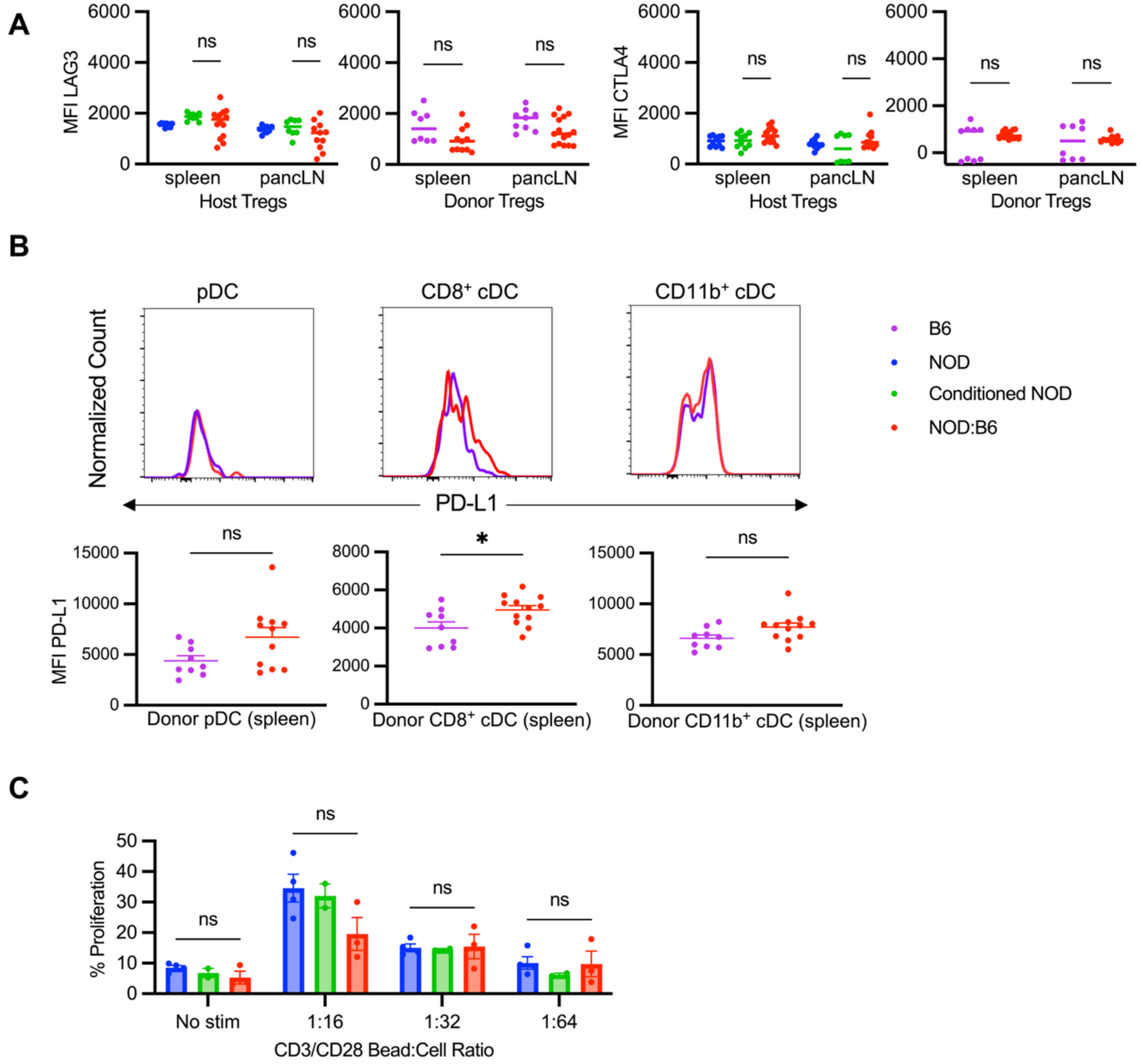


**Supplemental Figure 7. Peripheral tolerance mechanisms contribute to anergy of peripheral host effector cells.** (**A**) Median fluorescence intensity (MFI) of LAG3 and CTLA4 expressed by $CD45.1^{+}$ host Tregs in prediabetic NOD:B6 spleen and pancLN compared to NOD and conditioned NOD controls (*n* = 9-14). MFI of LAG3 and CTLA4 expressed by $CD45.2^{+}$ donor Tregs in NOD:B6 spleen and pancLN compared to B6 controls (*n* = 9-14). **(B)** Representative histogram and mean ± SEM of median fluorescence intensity (MFI) of PD-L1 expressed by $CD45.2^{+}$ donor plasmacytoid DCs (pDCs), $CD8^{+}$ conventional DCs (cDCs), and $CD11b^{+}$ cDCs in prediabetic NOD:B6 spleen compared to B6 controls(*n* = 9-10). (**C**) Proliferation of host $CD4^{+}$ splenic T cells from prediabetic NOD:B6 compared to NOD and conditioned NOD controls after isolation and incubation *in vitro* with CD3/CD28 stimulation beads at decreasing dilutions (1:16 to 1:64) compared to unstimulated. Data are represented with mean ± SEM. *P* values were calculated using Mann-Whitney tests $*P < 0.05$, $**P < 0.01$, $***P < 0.001$, $****P < 0.0001$; ns = not significant. DC = dendritic cell

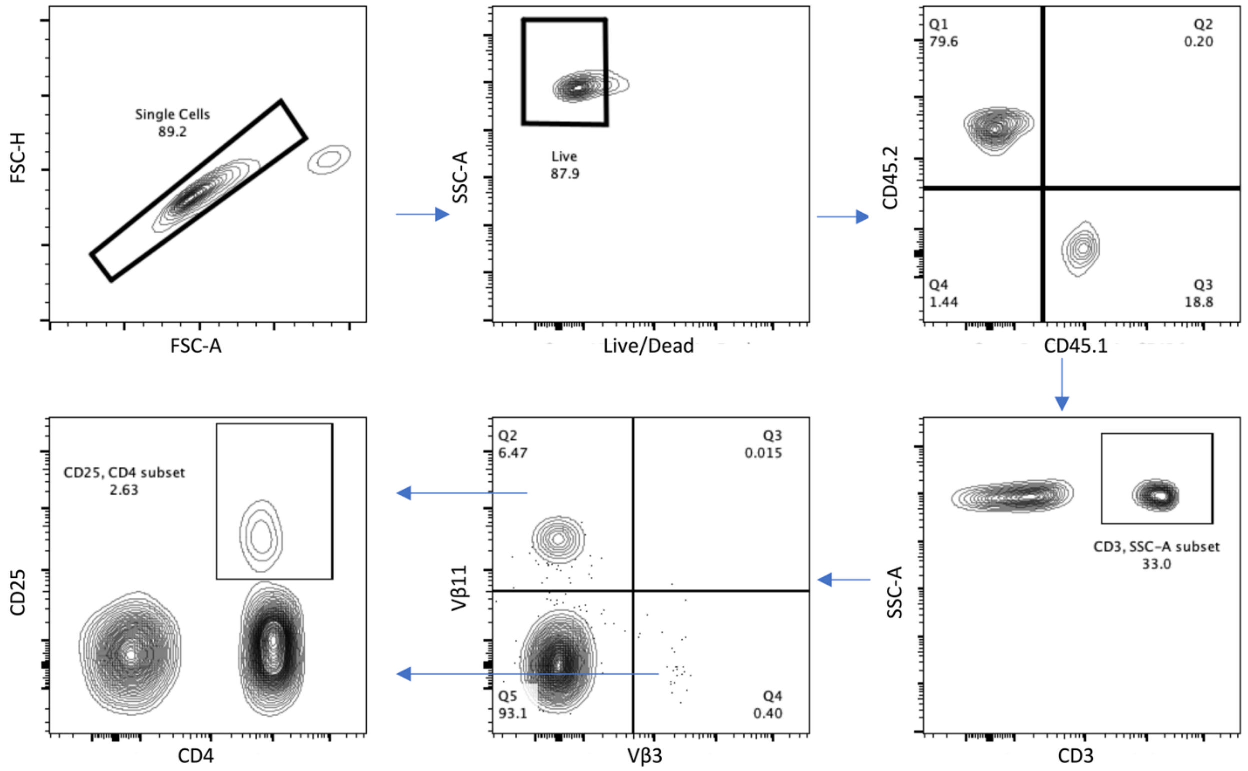


**Supplemental Figure 8. Gating strategy for Vβ subsets.** Lymphocytes were gated on FSC-A vs. SSC-A (not shown), then subset to single cell live donor ($CD45.2^+$) or host ($CD45.1^+$) $CD3^+$ T cells. These were further subset into $V\beta3^+$ and $V\beta11^+$ T cells. $CD4^+$ $CD25^+$ Tregs were gated on both $V\beta3^+$ and $V\beta11^+$ T cell subsets.

**Supplemental Table 1**

| **Primary Antibody** | **Dilution** | **Company** | **Catalog** |
|---|---|---|---|
| Rat α-Mouse CD3 | 1:200 | Biolegend | 100202 |
| Rat α-Mouse CD45 | 1:200 | Biolegend | 103102 |
| Guinea pig α-insulin | 1:500 | Dako | A0564 |
| Rabbit α -glucagon | 1:200 | ThermoFisher Scientific | PA5-88091 |
| **Secondary Antibody** | **Dilution** | **Company** | **Catalog** |
| AF647 Goat α-Guinea Pig | 1:100 | ThermoFisher Scientific | A-21450 |
| AF555 Donkey α-Rabbit | 1:100 | ThermoFisher Scientific | A-31572 |
| AF594 mouse α-Rat | 1:200 | Biolegend | 408207 |
| AF488 Mouse α-Rat | 1:200 | Biolegend | 408211 |
| AF594 Donkey α-Guinea Pig | 1:1000 | Millipore Sigma | SAB4600096 |
| AF488 Donkey α-Guinea Pig | 1:1000 | Millipore Sigma | SAB4600033 |
| **Conjugated Antibody** | **Dilution** | **Company** | **Catalog** |
| AF594 Rat α-Mouse CD45.2 | 1:100 | Biolegend | 109850 |
| AF488 Rat α-Mouse CD45.1 | 1:100 | Biolegend | 110717 |

**Supplemental Table 2**

| Antibody | Clone | Color | Catalog | Company |
|---|---|---|---|---|
| CD45.1 | A20 | PerCP-Cy-5.5 | 110728 | Biolegend |
| CD45.1 | A20 | BV685 | 110743 | Biolegend |
| CD45.2 | 104 | Pacific Blue | 109820 | Biolegend |
| CD3 | 17A2 | AF488 | 100210 | Biolegend |
| CD3 | 17A2 | AF700 | 100216 | Biolegend |
| CD3 | 17A2 | PE-Cy7 | 100220 | Biolegend |
| CD3 | 17A2 | FITC | 100204 | Biolegend |
| CD4 | RM4-4 | PE | 116006 | Biolegend |
| CD4 | RM4-4 | BV421 | 100563 | Biolegend |
| CD8 | 53-6.7 | BV510 | 100752 | Biolegend |
| CD8 | 53-6.7 | FITC | 100705 | Biolegend |
| CD11b | M1/70 | BV605 | 101257 | Biolegend |
| CD11b | M1/70 | FITC | 101206 | Biolegend |
| CD25 | PC61.5 | PE-Cy5 | 102010 | Biolegend |
| CD19 | 6D5 | PE-Cy7 | 115520 | Biolegend |
| CD49b | DX5 | APC | 108910 | Biolegend |
| Gr-1 | R B6-8C5 | FITC | 108406 | Biolegend |
| TER-119 | TER-119 | FITC | 116206 | Biolegend |
| CD11c | N418 | BV421 | 117329 | Biolegend |
| CD172a (SIRPa) | P84 | APC | 144013 | Biolegend |

| | | | | |
|---|---|---|---|---|
| CD317 (PDCA-1) | 927 | PE | 127009 | Biolegend |
| MHCII (I-A/I-E) | M5/114.15.2 | PE-Cy7 | 107629 | Biolegend |
| CD274 (PDL1) | 10F.9G2 | PE/Dazzle 594 | 124323 | Biolegend |
| B220 | RA3-6B2 | FITC | 103206 | Biolegend |
| CD62L | MEL-14 | PE-Cy5 | 104410 | Biolegend |
| CD44 | IM7 | BV605 | 103047 | Biolegend |
| CD304 (Nrp1) | 3E12 | PE | 145203 | Biolegend |
| CD73 | TY/11.8 | PE/Dazzle 594 | 127234 | Biolegend |
| FR4 | 12A5 | PE-Cy7 | 125012 | Biolegend |
| Helios | 22F6 | AF488 | 137213 | Biolegend |
| FOXP3 | 150D | AF647 | 320013 | Biolegend |
| CD39 | Duha59 | PE/Fire 640 | 143818 | Biolegend |
| CD19 | 6D5 | AF700 | 115528 | Biolegend |
| TCR-beta | H57-597 | AF700 | 109224 | Biolegend |
| ICOS (CD278) | C398.4A | BV650 | 313549 | Biolegend |
| LAG3 (CD223) | C9B7W | BV711 | 125243 | Biolegend |
| CTLA4 (CD152) | UC10-4B9 | BV421 | 106311 | Biolegend |
| PD1 | 29F.1A12 | BV750 | 135263 | Biolegend |
| CD25 | 3C7 | PE-Cy7 | 101915 | Biolegend |
| CD4 | GK1.5 | AF700 | 100429 | Biolegend |
| CD4 | GK1.5 | Pacific Blue | 116008 | Biolegend |
| CD117 | 2B8 | APC | 17-1171-82 | ThermoFisher Scientific |
| Sca-1 | D7 | PE-Cy7 | 25-5981-81 | ThermoFisher Scientific |

| | | | | |
|---|---|---|---|---|
| TCR Vβ11 | RR3-15 | PE | 139004 | Biolegend |
| TCR Vβ3 | REA646 | APC | 130-109-895 | Miltenyi Biotec |
